# Related PP2C phosphatases Pic3 and Pic12 negatively regulate immunity in tomato to *Pseudomonas syringae*

**DOI:** 10.1101/2024.02.08.579555

**Authors:** Fan Xia, Ning Zhang, Renee E. Smith, Joydeep Chakraborty, Guy Sobol, Xuemei Tang, Zhangjun Fei, Guido Sessa, Gregory B. Martin

## Abstract

Type 2C protein phosphatases (PP2Cs) constitute a large family in most plant species but relatively few of them have been implicated in immunity. To identify and characterize PP2C phosphatases that affect tomato immunity, we used CRISPR/Cas9 to generate loss-of-function mutations in 11 PP2C-encoding genes whose expression is altered in response to immune elicitors or pathogens. We report that two closely related PP2C phosphatases, Pic3 and Pic12, are involved in regulating resistance to the bacterial pathogen *Pseudomonas syringae* pv. *tomato* (*Pst*). Loss-of-function mutations in *Pic3* lead to enhanced resistance to *Pst* in older but not younger leaves, whereas such mutations in *Pic12* resulted in enhanced resistance in both older and younger leaves. Overexpression of Pic3 and Pic12 proteins in leaves of *Nicotiana benthamiana* inhibited resistance to *Pst* and this effect was dependent on Pic3/12 phosphatase activity and an N-terminal palmitoylation motif associated with localization to the cell periphery. Pic3 but not Pic12 had a slight negative effect on flagellin-associated reactive oxygen species generation, although their involvement in the response to *Pst* appeared independent of flagellin. RNA-sequencing analysis of Rio Grande (RG)-PtoR wild-type plants and two independent RG-pic3 mutants revealed that the enhanced disease resistance in RG-pic3 older leaves is associated with increased transcript abundance of multiple defense related genes. RG-pic3/RG-pic12 double mutant plants exhibited stronger disease resistance than RG-pic3 or RG-pic12 single mutants. Together, our results reveal that Pic3 and Pic12 negatively regulate tomato immunity in an additive manner through flagellin-independent pathways.

## Introduction

Plants use a two-layered innate immune system to protect themselves against pathogen attack. The first layer uses cell surface-associated pattern recognition receptor (PRR) proteins to detect pathogen associated molecular patterns (PAMPs) to activate PRR-triggered immunity (PTI). Successful bacterial pathogens translocate effector proteins into plant cells through the type III secretion system (T3SS) and, among several activities, target various host components to inhibit PTI. The second layer of defense deploys intracellular nucleotide-binding, leucine-rich repeat (NLR) family proteins to sense pathogen effectors and initiate effector triggered immunity (ETI) (Jones and Dangl, 2006). Despite the differences in recognition mechanisms, PTI and ETI both elicit some common host responses including reactive oxygen species (ROS) burst and calcium fluxes, activation of mitogen activated protein kinase (MAPK), and transcriptional reprograming and ultimately result in the restriction of pathogen multiplication (Zhang et al., 2020; Wang and Luan, 2023).

A well-characterized plant PRR, Flagellin Sensing 2 (Fls2), is a receptor kinase conserved in higher plants that binds a 22 amino acid epitope, flg22, of the bacterial flagellin protein (Sun et al., 2013; Cheng, Xiao and Xiong, 2020). In tomato and other solanaceous plants, a distinct 28 amino acid epitope, flgII-28, also derived from flagellin, is recognized by the receptor kinase Flagellin Sensing 3 (Fls3) (Clarke et al., 2013; Hind et al., 2016). Fls2 and Fls3 each form a complex with co-receptor BRASSINOSTEROID INSENSITIVE 1–associated kinase 1 (BAK1) to activate downstream defense responses (Chinchilla et al., 2007; Hind et al., 2016). Fls2-and Fls3-mediated PTI plays an important role in tomato resistance to bacterial pathogens (Rosli et al., 2013; Roberts et al., 2020). Consequently, either the deletion of bacterial flagellin-encoding gene *fliC* or loss-of-function mutations in both Fls2 and Fls3 significantly compromise tomato resistance against *Pseudomonas syringae* pv. *tomato* (*Pst*) DC3000 (Roberts et al., 2020; Zhang et al., 2020). The cold shock protein (CSP)-derived bacterial PAMP, csp22, is recognized by the receptor kinase cold shock protein receptor (CORE) in tomato and loss-of-function mutations in the *CORE* gene in tomato lead to enhanced susceptibility to *Pst* (Danielle et al., 2024).

The tomato disease resistance gene *Pto* encodes a protein kinase that physically interacts with T3SS effectors AvrPto and AvrPtoB (Tang et al., 1996; Kim, Lin and Martin, 2002). However, to trigger ETI, Pto interacts and forms a complex with the NLR protein Prf (Ntoukakis et al., 2013). A number of signaling components acting downstream of Pto/Prf have been identified, including NLR required for cell death (Nrc2 and Nrc3), 14-3-3 proteins Tomato Fourteen-Three-three (Tft1, Tft3 and Tft7), mitogen-activated protein (MAP) kinase cascade components MAP Kinase Kinase Kinase (MAPKKKα), MAP kinase kinases (MKK1 and MKK2) and MAP Kinases (wound-inducible protein kinase (WIPK) and NTF6)), and M3Kα-interacting protein 1 (Mai1) (Ekengren et al., 2003; Mucyn et al., 2006; Oh, Pedley and Martin, 2010; Oh and Martin, 2011; Roberts et al., 2019; Sheikh et al., 2023; Zhang et al., 2024). Loss of Pto/Prf or of certain downstream components disrupts Pto/Prf mediated ETI, resulting in compromised resistance against *Pst* stains carrying AvrPto or AvrPtoB.

PTI and ETI signaling involve many protein kinases which rely on phosphorylation for promoting signal transduction (Ngou, Ding and Jones, 2022). In contrast, protein phosphatases counteract kinase function by dephosphorylating their substrates. Protein phosphatases together with kinases act as molecular switches and maintain balanced and tightly regulated defense responses (Erickson et al., 2022). For instance, overexpression of kinase associated protein phosphatase (KAPP) in Arabidopsis inhibits the binding of FLS2 with flagellin and results in reduced flagellin sensitivity (Gomez-Gomez, Bauer and Boller, 2001). A heterotrimeric protein Ser/Thr phosphatase type 2A (PP2A), comprised of several subunits, has been shown to regulate Arabidopsis immunity through modulating the phosphorylation status of FLS2 co-receptor BAK1 (Segonzac et al., 2014). Additionally, silencing a catalytic subunit of PP2A (PP2Ac) in *Nicotiana benthamiana* reduces PP2A activity and enhances the hypersensitive response (HR) induced by avirulent *Pst* (He et al., 2004).

In Arabidopsis, there are about 1,125 protein kinases but only around 150 protein phosphatases (Bheri et al., 2021). Notably, much less is known about protein phosphatases than kinases in plant defense, partly because forward genetic screening of defense-related mutants in the past has rarely identified protein phosphatases (He et al., 2004). Based on protein sequence and catalytic mechanism, plant protein phosphatases are divided into four families: phosphoprotein phosphatase (PPP), protein phosphatase 2C (PP2C, also named Mg^2+^/Mn^2+^-dependent protein phosphatase (PPM)), phosphotyrosine phosphatase (PTP) and Asp-dependent phosphatase (Uhrig, Labandera and Moorhead, 2013). The PPP and PP2C family members are serine/threonine protein phosphatases that dephosphorylate phospho-serine and phospho-threonine but the sequences of PP2Cs are distinct from those of PPP members (Uhrig, Labandera and Moorhead, 2013). Significant attention has been given to PP2Cs in recent years as many of them are found to regulate biotic and abiotic stresses (Schweighofer, Hirt and Meskiene, 2004; Sobol et al., 2022). For example, several PP2Cs are regulators of abscisic acid (ABA) signaling, which plays an important role in abiotic stress responses and other physiological processes (Chen et al., 2020). In Arabidopsis, several PP2C phosphatases have been implicated in dampening immunity by dephosphorylating defense signaling components. These include Highly ABA-Induced1 (HAI1) and Arabidopsis PP2C-type phosphatase 1 (AP2C1) that dephosphorylate MAPKs and PP2C38 that dephosphorylates Botrytis-Induced Kinase 1 (BIK1), POLTERGEIST-LIKE 4 (PLL4) that mediates dephosphorylation of FLS2 and the Elongation Factor Tu Receptor (EFR), and chitin elicitor receptor kinase1 (CERK1)-interacting protein phosphatase 1 (CIPP1) that dephosphorylates chitin co-receptor CERK1 (Schweighofer et al., 2007; Couto et al., 2016; Mine et al., 2017; Liu et al., 2018; DeFalco et al., 2022). In rice, PP2C phosphatase XA21 binding protein 15 (XB15) mediates receptor kinase XA21 dephosphorylation and the loss of XB15 activates cell death, increases defense-related gene expression, and enhances XA21-mediated resistance (Park et al., 2008).

Most research about plant PP2Cs has been performed in Arabidopsis and to a lesser extent in rice. Little is known about the role of these phosphatases in the tomato immune response. Compared to Arabidopsis and rice, which have 80 and 76 PP2C phosphatases, respectively, the tomato (*Solanum lycopersicum*) genome encodes 97 PP2C phosphatases (Xue et al., 2008; Fuchs et al., 2013; Qiu et al., 2022; Sobol et al., 2022). Many tomato PP2Cs share high sequence similarity with their Arabidopsis and rice counterparts, suggesting they might play a conserved role in these species. We previously reported that the tomato PP2C phosphatase Pic1 negatively modulates ROS production in *N. benthamiana* by dephosphorylating a key PTI related kinase, Pti1b (Schwizer et al., 2017; Giska and Martin, 2019). By examining transcriptomic data, we subsequently found that transcript abundances of *Pic1* and 12 other tomato PP2C genes were significantly increased or decreased upon PAMP and pathogen treatments, suggesting they might have a role in immunity (Sobol et al., 2022). We now refer to these as <u>P</u>P2C immunity-associated <u>c</u>andidate (Pic) proteins. Here we used CRISPR/Cas9 technology to generate mutations in a collection of *Pic* genes and report that two closely related Pic phosphatases negatively regulate tomato immunity.

## Results

### A screen of PP2C immunity candidate (Pic) mutant lines reveals that Pic3 negatively regulates resistance to *P. syringae* pv. *tomato* in older but not younger leaves

To assess the role of Pic phosphatases in the tomato defense response, we first generated gene-specific guide RNAs (gRNAs) and used CRISPR/Cas9 technology to induce loss-of-function mutations in a set of eight previously identified PP2C-encoding genes and, in two cases (*Pic7* and *Pic10*), an additional two closely-related genes (Sobol et al., 2022) (**Figure 1A** and **Supplemental Figure S1**). Agrobacterium-mediated stable transformation with the 10 gRNA constructs was performed using tomato cultivar Rio-Grande PtoR (RG-PtoR, wildtype), which expresses the *Pto/Prf* genes and recognizes race 0 *Pseudomonas syringae* pv. *tomato (Pst)* strains (Pedley and Martin, 2003). Cas9-mediated target gene disruption resulted in deletions and insertions near the 5’ end of the *Pic* genes, which caused open-reading frameshifts and introduced a premature stop codon (except for *Pic11*), leading to predicted loss-of-function mutations (**Figure 1A** and **Supplemental Figures S1** and **S2A**). For most *Pic* genes, we obtained multiple transgenic lines (referred to as RG-pic lines) and chose 1 or 2 independent lines for each gene for further study. The overall morphology and growth of 4-week-old RG-pic mutant plants was generally indistinguishable from wild-type plants (**Figure 1A** and **Supplemental Figure S3A**).

**Figure 1.**
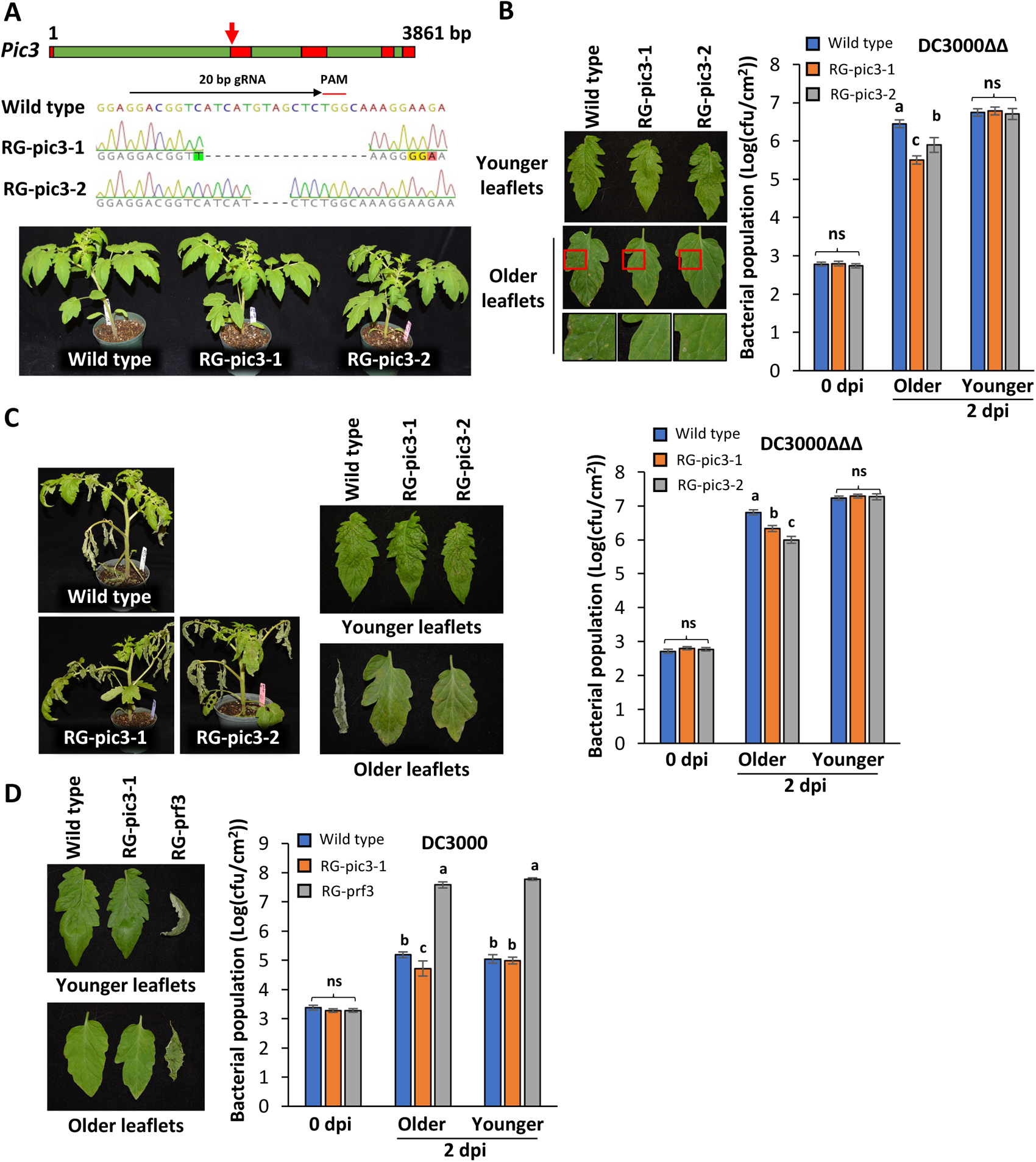
Tomato loss-of-function RG-pic3 mutants show increased disease resistance. **A)** Top panel, *Pic3* gene structure including exons (red) and introns (green), arrow points to CRISPR/Cas9 gRNA target site; Middle panel, sequencing chromatogram of two missense mutant alleles generated by CRISPR/Cas9; Bottom panel, images of 4-week-old wild-type (Rio Grande-PtoR; RG-PtoR) and RG-pic3 mutant plants. **B)** Wild-type and RG-pic3 plants were vacuum-infiltrated with 5×10^4^ cfu/mL DC3000Δ*avrPto*Δ*avrPtoB* (DC3000ΔΔ) and photographed at 4 days post-inoculation (dpi); a section of the older leaflets was enlarged for better visualization. **C)** Wild-type and RG-pic3 plants were vacuum-infiltrate with 5×10^4^ cfu/mL DC3000Δ*avrPto*Δ*avrPtoB*Δ*fliC* (DC3000ΔΔΔ) and photographed at 4 dpi. **D)** Wild-type, RG-pic3 and RG-prf3 plants were vacuum-infiltrated with 2×10^5^ cfu/mL DC3000 and photographed at 4 dpi. ‘Older’ refers to 1^st^ or 2^nd^ true leaf and ‘younger’ refers to 4^th^ or 5^th^ true leaf. Bacterial populations in all panels were measured at 0 and 2 dpi. Bars show means ± standard deviation, n = 3 or 4. cfu, colony-forming unit. Different letters indicate significant differences based on a one-way ANOVA followed by Tukey’s Honest Significant Difference post hoc test (P < 0.05). ns, no significant difference. All inoculation experiments were repeated three times with similar results.

To test if these *Pic* genes are involved in tomato defense responses to *Pst*, we inoculated each RG-pic line with three different *Pst* DC3000 strains. DC3000Δ*avrPto*Δ*avrPtoB* (DC3000ΔΔ) carries deletions in two genes encoding type III effectors that are recognized by Pto/Prf (Lin and Martin, 2005). Thus, it does not trigger ETI and can be used to test PTI-based responses in RG-PtoR plants. The DC3000Δ*avrPto*Δ*avrPtoB*Δ*fliC* (DC3000ΔΔΔ) strain, has an additional deletion of the *fliC* gene encoding the flagellin protein. Since flagellin-associated peptides, flg22 and flgII-28, are the major *Pst* PAMPs recognized by tomato (via the Fls2 and Fls3 receptors, respectively), this strain can reveal potential non-flagellin-based basal defense responses in tomato (Rosli et al., 2013; Hind et al., 2016; Roberts et al., 2020). The wild-type DC3000 strain is used to test for AvrPto-and AvrPtoB-activated ETI responses (Zhang et al., 2020). Our screening of the RG-pic mutant lines showed that loss-of-function mutations in nine of the *Pic* genes (two genes each in the RG-pic7 and RG-pic10 lines) did not significantly affect the tomato response to any of the DC3000 strains, as bacterial populations in these RG-pic mutants were not statistically different from wild-type plants (**Supplemental Figures S3B, S3C** and **S3D).**

Two independent lines were identified that had predicted loss-of-function mutations in the *Pic3* gene (Solyc01g105280). Line RG-pic3-1 has a 17-bp deletion and 4-bp substitutions in exon 2 and line RG-pic3-2 has a 4-bp deletion in exon 2 (**Figure 1A**). In both cases, the mutations are predicted to result in truncated proteins lacking the PP2C domain (**Supplemental Figure S2A**). Inoculation of the lines with DC3000ΔΔ resulted in fewer bacterial specks in the older leaves of the RG-pic3 lines than on wild-type plants at 4 days after inoculation (**Figure 1B**). In addition, lower bacterial populations were observed in older leaves of the RG-pic3 lines than in wild-type plants at 2 days post-inoculation (dpi; **Figure 1B**). Interestingly, DC3000ΔΔ disease symptoms and populations in younger leaves of the RG-pic3 mutant lines were not different from those of wild-type plants (**Figure 1B**).

When inoculated with DC3000ΔΔΔ, we observed less severe disease symptoms and markedly reduced bacterial population in older leaves of both RG-pic3 lines compared to wild-type plants, while DC3000ΔΔΔ disease symptoms and populations in younger leaves were not different between wild-type and RG-pic3 plants (**Figure 1C**). These results suggest that Pic3 negatively regulate PTI and/or other basal defenses, and this regulation is at least partially independent of flg22 and flgII-28. Since both RG-pic3-1 and RG-pic3-2 showed similar phenotypes in these initial experiments we used just the RG-pic3-1 line in some of the subsequent experiments.

As expected, DC3000 at a relatively low titer (2x10^5^ cfu/mL) activates ETI and did not cause symptoms on wild-type RG-PtoR plants, but bacterial populations in this line still slightly increased at 2 dpi. The RG-pic3 plants supported slightly less DC3000 growth than wild-type plants in the older but not younger leaves (**Figure 1D**). However, RG-pic3 plants inoculated with a higher titer of DC3000 (1x10^7^ cfu/mL) developed a hypersensitive response that was indistinguishable from wild-type RG-PtoR plants (**Supplemental Figure S4**), indicating that Pic3 does not play a major role in regulating ETI. Finally, we observed that older leaves of 8-week-old plants of the two RG-pic3 lines started to show early senescence while those of wild-type plants did not (**Supplemental Figure S2B**), suggesting an autoimmune response or Pic3-mediated regulation of additional processes in addition to defense.

Pic3 protein belongs to clade D or clade F of PP2Cs, depending on the phylogenetic analysis (Fuchs et al., 2013; Qiu et al., 2022; Sobol et al., 2022). Several clade A PP2Cs have been reported to regulate ABA signaling, which in some cases is involved in senescence development (Tischer et al., 2017; Chen et al., 2020). To test if the enhanced disease resistance seen in RG-pic3 plants might be associated with an ABA response, we examined the expression of two well-known ABA-pathway reporter genes in RG-pic3 plants after exogenous ABA treatment (Ma et al., 2018; Lei et al., 2023). The expression levels of these two genes, *AREB1* and *SAG113*, were not different in RG-pic3 plants compared to wild-type plants (**Supplemental Figures S5A** and **S5B**), suggesting Pic3-associated defense is unlikely to be associated with an ABA response.

### Pic3 phosphatase activity and its palmitoylation motif are required for its negative regulation of the tomato immune response

*Pic3* encodes a PP2C protein with a single PP2C catalytic domain but no known regulatory domain at the N and C terminus (Sobol et al., 2022). The catalytic domain contains conserved aspartic acid residues that play a role in binding metal ions such as Mn^2+/^Mg^2+^ (**Supplemental Figure S6**), and several studies (Jackson, Fjeld and Denu, 2003; Conner et al., 2006; Couto et al., 2016; Giska and Martin, 2019) reported that substitutions from aspartic acid (D) to asparagine (N) at these highly conserved sites abolishes phosphatase activities. To test the possible requirement of Pic3 phosphatase activity in defense, we generated a construct carrying a variant, Pic3NN, with substitutions at the two conserved aspartic residues (D71N and D233N; **Figure 2A**). Using the ‘agromonas’ approach (Buscaill et al., 2021), we transiently expressed Pic3, Pic3NN and a YFP control for 2 days in *N. benthamiana* leaves and then inoculated the plants with DC3000*ΔhopQ1-1,* which lacks the T3SS effector gene *hopQ1-1* that can be detected by the NLR protein Recognition of XopQ 1 (Roq1), thus making DC3000*ΔhopQ1-1* virulent in *N. benthamiana;* (Wei et al., 2007; Schultink et al., 2017). Compared to the YFP control, transient expression of wild-type Pic3 significantly increased the DC3000*ΔhopQ1-1* population in *N. benthamiana,* consistent with Pic3 playing a negative role in resistance to DC3000*ΔhopQ1-1* (**Figure 2B**). In contrast, bacterial abundance in leaves expressing Pic3NN were not statistically different from that in plants expressing YFP (**Figure 2B**), indicating that phosphatase activity plays an important role in Pic3 function in defense. Consistent with the observation that Pic3-modulated defense is flagellin independent, when inoculated with a flagellin deficient strain, DC3000*ΔhopQ1-1ΔfliC*, *N. benthamiana* plants expressing wild-type Pic3 protein also supported significantly higher bacterial populations than plants expressing Pic3NN or YFP (**Figure 2C**). Immunoblot analysis confirmed that the Pic3 and Pic3NN proteins were expressed in leaves at similar levels (**Figure 2D**).

**Figure 2.**
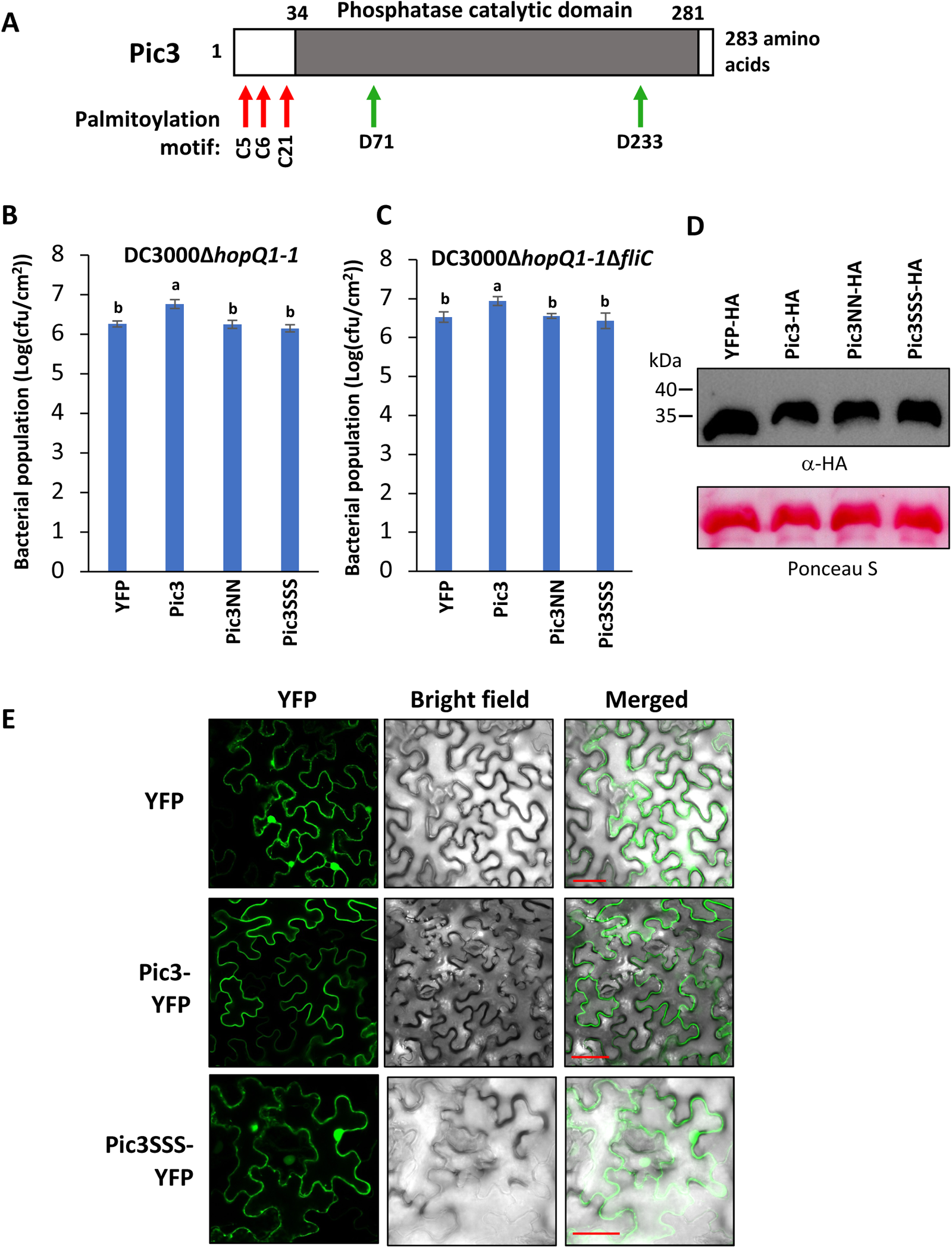
Phosphatase activity and palmitoylation motif are required for Pic3 function in defense and palmitoylation motif affects subcellular localization of Pic3. **A)** Schematic of Pic3 protein showing the three predicted palmitoylation sites (red arrows) at the N terminus and two conserved aspartic acid residues (green arrows) involved in metal binding in the PP2C catalytic domain. **B a**nd **C)** Leaves of 5-week-old *N. benthamiana* plants were syringe-infiltrated with Agrobacterium (1D1249) strains (OD_600_ = 0.6) carrying a binary expression vector (pGWB414) expressing 3× HA-tagged wild-type Pic3, a phosphatase-inactive variant Pic3NN (D71N and D233N substitutions), or a Pic3SSS variant with substitutions in the palmitoylation motif (C5S, C6S and C21S) or the YFP control. Two days later, the same agroinfiltrated spots were syringe-infiltrated with 5×10^4^ cfu /mL DC3000*ΔhopQ1-1* (B) or 5×10^4^ cfu/mL DC3000*ΔhopQ1-1ΔfliC* (C)*. Pst* DC3000 populations were measured at 2 days after the second infiltration. Bars show means ± standard deviation, n = 4. Different letters indicate significant differences based on a one-way ANOVA followed by Tukey’s Honest Significant Difference post hoc test (P < 0.05). Experiments were repeated three times with similar results. **D)** Western blotting analysis with an anti-HA antibody confirmed that proteins tested in (B and C) were expressed. *N. benthamiana* protein samples were collected at 40 h after initial agrobacterium infiltration. **E)** Wild-type Pic3 or the Pic3SSS variant protein fused with YFP (pGWB541 vector) were transiently expressed in *N. benthamiana* leaves via Agrobacterium (GV3101) for 40 h and examined under a confocal microscope. YFP fluorescence was excited by 485 nm laser. The pGWB541 vector (CmR, ccdB removed) expressing YFP was used as a control. Scale bar, 50 μm. Experiments were repeated twice with similar results.

We next discovered, using GS-Palm (Ning et al., 2021), that three cysteine residues (C5, C6 and C21) at the Pic3 N-terminus constitute a predicted palmitoylation motif. Palmitoylation is the covalent addition of fatty acids to a cysteine residue, which often promotes association of proteins with membranes (Lin, 2021). As our disease assays indicate that Pic3 could negatively regulate a PTI response, we hypothesized that Pic3 might associate with the plasma membrane to modulate early PTI-related signaling events. To test the possible requirement of Pic3 palmitoylation in defense, we generated a construct carrying a Pic3 variant with a disrupted palmitoylation motif (Pic3SSS; with C5S, C6S and C21S substitutions) and tested it using the agromonas approach described above (**Figures 2A, 2B** and **2C**). We observed that bacterial abundances in Pic3SSS-expressing plants were not statistically different from those in plants expressing YFP or Pic3NN (**Figures 2B** and **2C**), suggesting that palmitoylation is required for Pic3 function in defense. Immunoblot analysis confirmed that the Pic3 and Pic3SSS proteins were expressed at similar levels (**Figure 2D**).

To test if the palmitoylation motif affects Pic3 subcellular localization, we fused a YFP tag to the C-terminus of Pic3 and Pic3SSS and examined Pic3-and Pic3SSS-YFP fusion protein localization with a confocal microscope. As expected, we found that fluorescence from the YFP protein control was present in both the cytosol and nucleus. Fluorescence from the Pic3-YFP protein was noticeably associated with the cell periphery, whereas a significant portion of Pic3SSS-YFP fluorescence appeared in the nucleus (**Figure 2E**). Immunoblot analysis confirmed that the Pic3-YFP and Pic3SSS-YFP proteins were expressed at similar levels (**Supplemental Figure S7A**). These observations support the hypothesis that palmitoylation of the Pic3 protein affects its subcellular localization. We also investigated, but did not find, an effect on Pic3 protein localization associated with inoculation by DC30000*ΔhopQ1-1* or exposure to the PAMP flg22 (**Supplemental Figure S7B**).

### Pic3 does not affect MAPK activation but has a minor role in negatively regulating flgII-28 mediated ROS production

MAPK activation and ROS production are two PTI-associated responses that occur in tomato after the recognition of flg22 or flgII-28 by Fls2 or Fls3, respectively (Roberts et al., 2020; Zhang et al., 2020). Although we showed that Pic3-modulated defense against *Pst* can be independent of flagellin, it remained possible that certain responses controlled by Fls2 and/or Fls3 might be regulated by this phosphatase. We therefore monitored MAPK activation in wild-type and RG-pic3 plants after exposure to flg22 or flgII-28 (Chinchilla et al., 2006; Hind et al., 2016; Roberts et al., 2020). We also included in our MAPK assay another PAMP, csp22 peptide, which is derived from a cold shock protein, and recognized by the CORE receptor (Wang et al., 2016; Danielle et al., 2024) to see if it was involved in the flagellin-independent response we observed earlier. MAPKs were activated similarly in both older and younger leaves in wild-type and RG-pic3 plants with all three different peptide treatments (**Figure 3A**). Other unknown PAMPs might also induce PTI and trigger MAPK activation in tomato. To test if MAPK activation due to these PAMPs is affected by Pic3, we used boiled *Pst* cell lysates from DC3000ΔΔ and DC3000ΔΔΔ as a source of other unknown PAMPs to induce MAPK. However, we again detected no significant difference between RG-pic3 and wild-type plants in MAPK activation in response to these lysates (**Supplemental Figure S8**).

**Figure 3.**
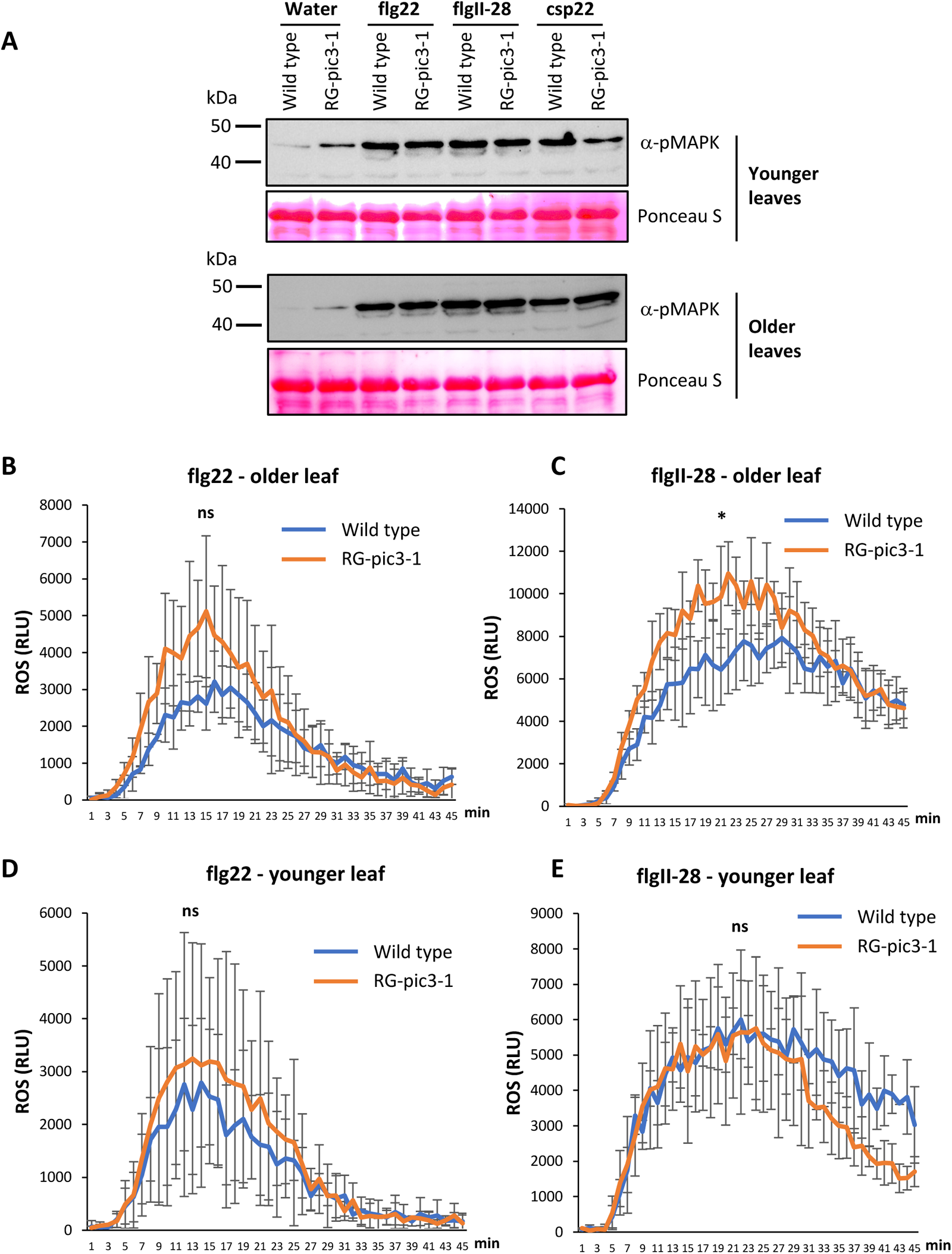
MAPK activation and ROS production in RG-pic3 mutant plants. **A)** MAPK activation in wild-type (RG-PtoR) and RG-pic3 mutant plants. Water, 10 nM flg22, 25 nM flgII-28 or 50 nM csp22 were applied to detached leaf discs for 10 min and proteins were separated on 12% SDS-PAGE gel and detected by an anti-pMAPK antibody. **B** and **C)** Time course of ROS production in older leaves of wild-type and RG-pic3 mutant plants treated with 100 nM flg22 or 100 nM flgII-28. **D** and **E)** Time course of ROS production in younger leaves of wild-type and RG-pic3 mutant plants treated with 100 nM flg22 or 100 nM flgII-28. ROS production was measured as relative light units (RLUs) for 45 min. Bars show means ± standard deviation, n = 3 plants. An asterisk indicates significant differences between wild-type and RG-pic3 plants at peak readout based on Student’s t-test (P < 0.05), ns, no significant difference. ‘Older’ refers to 1^st^ or 2^nd^ true leaf and ‘younger’ refers to 4^th^ or 5^th^ true leaf. Experiments were repeated three times with similar results.

To test if Pic3 affects Fls2-or Fls3-mediated ROS production, we measured ROS production in older and younger leaves of RG-pic3 and wild-type plants after flg22 or flgII-28 treatment. We observed no statistically significant increase in flg22-induced ROS in older leaves of RG-pic3 compared to wild-type plants (**Figure 3B**). However, after flgII-28 treatment, older leaves of RG-pic3 plants did consistently show higher ROS production than those of wild-type plants (**Figure 3C**). We observed no difference in ROS production in younger leaves between RG-pic3 and wild-type plants in response to either flg22 or flgII-28 (**Figures 3D** and **3E**). Overall, these results suggest that Pic3 has a slight negative effect on flgII-28-activated ROS production but otherwise does not have a detectable effect on two well-characterized PTI responses.

### Transcript abundance of defense-related genes in RG-pic3 plants is increased in older leaves compared to wild-type plants

*Pst* is known to activate significant transcriptional reprograming in tomato, which is associated with restriction of bacterial multiplication in leaves (Rosli et al., 2013; Pombo et al., 2014; Zhang et al., 2022). To investigate how Pic3 might affect the plant immune response independent of flagellin, we vacuum inoculated wild-type plants and the two RG-pic3 lines with DC3000ΔΔΔ and collected RNA samples from older leaves and younger leaves separately at 6 h post-inoculation (hpi) and carried out an RNA-sequencing (RNA-Seq) analysis. This analysis revealed that substantially more differently expressed genes (DEGs) were found in older than younger leaves of RG-pic3 compared to wild-type plants (**Figure 4A**). As we have done previously to simplify the analysis (Rosli et al., 2013; Pombo et al., 2014), we focused on genes with an FPKM (fragments per kilobase of transcript per million mapped fragments) of ≥3 in RG-pic3 (for up-regulated DEGs) or wild-type plants (for down-regulated DEGs), and a fold-change of ≥2 and an FDR (false discovery rate) of < 0.05. Applying these parameters to older leaves, we found that transcript abundance of 1,123 genes was greater in RG-pic3 plants compared to wild-type plants and transcript abundance of 387 genes was less in RG-pic3 compared to wild-type plants (**Figure 4A**). In contrast, in younger leaves there were only 13 genes with higher transcript abundance in RG-pic3 compared to wild-type plants and just two genes with lower transcript abundance in RG-pic3 compared to wild-type plants. These results are consistent with the observation that RG-pic3 plants have stronger resistance to DC3000ΔΔΔ than wild type plants in older but not in younger leaves (**Figure 1C**).

**Figure 4.**
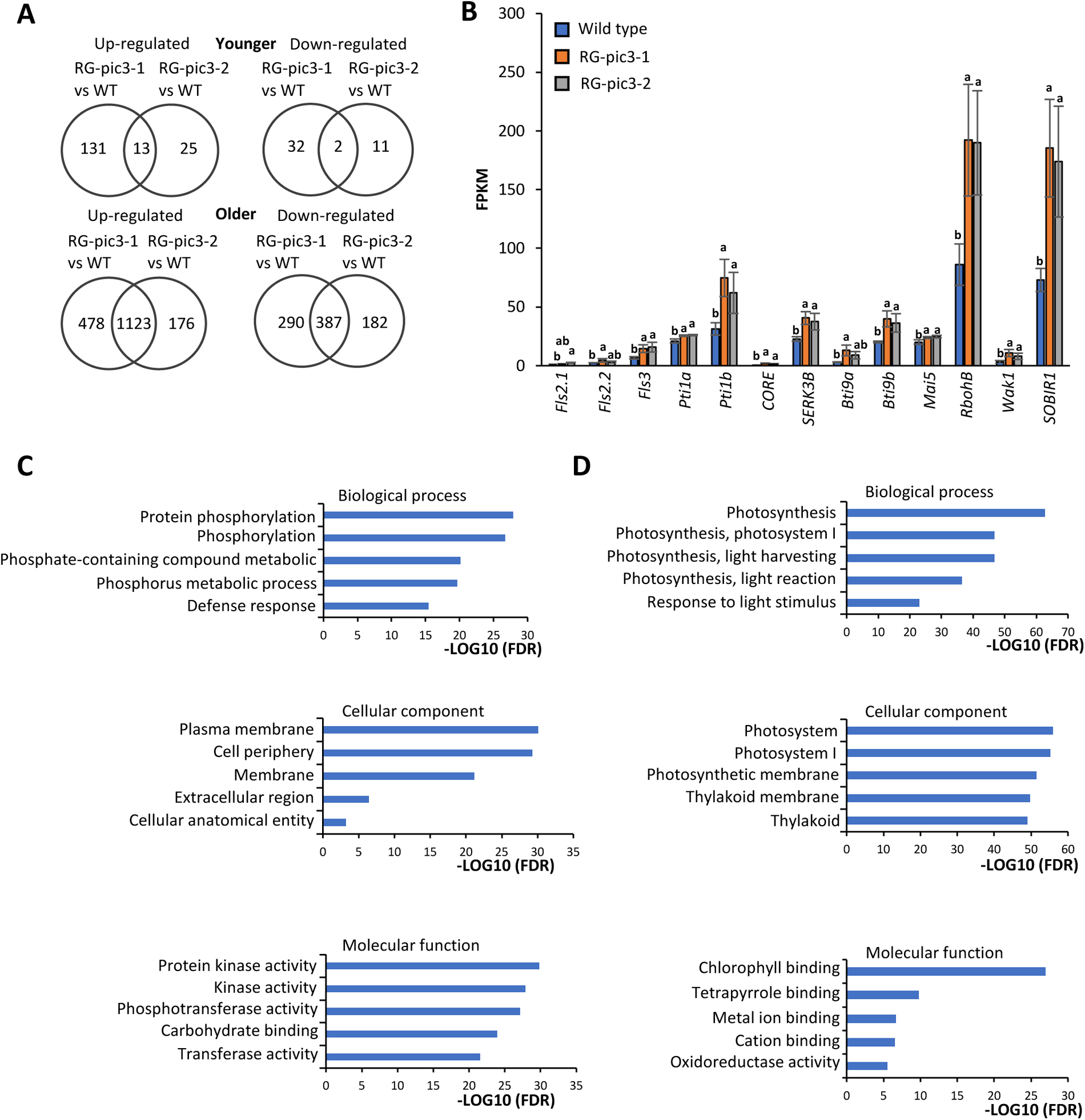
Enhanced disease resistance in RG-pic3 older leaves is associated with higher defense gene expression. **A)** Venn diagrams showing number of differentially-expressed genes (DEGs) in RG-pic3 and wild-type plants identified through RNA-Seq. Fold change >= 2, P < 0.05 and FPKM (fragments per kilobase of transcript per million mapped fragments) >= 3 in at least one genotype. ‘Older’ refers to 1^st^ or 2^nd^ true leaf and ‘younger’ refers to 4^th^ or 5^th^ true leaf. **B)** Expression levels of several known PTI-associated genes in wild-type and RG-pic3 plants older leaves at 6 h after DC3000Δ*avrPto*Δ*avrPtoB*Δ*fliC* inoculation (derived from RNA-Seq data). **C)** GO-enriched terms analysis focused on up-regulated DEGs in older leaves of RG-pic3 compared to that of wild-type plants. Only top 5 terms with lowest FDR (false discovery rate) are shown. **D)** GO-enriched terms analysis focused on down-regulated DEGs in older leaves of RG-pic3 compared to that of wild-type plants. Only the top 5 terms with lowest FDR are shown. See Supplemental Table S1 for full list of significantly enriched GO terms.

Transcript abundance of several genes associated with PTI was higher in older leaves of RG-pic3 compared to wild-type (**Figure 4B**), and it is possible that the increased expression of these genes contributes to the enhanced resistance in RG-pic3 older leaves. To gain broader insight into how the 1,123 upregulated DEGs might impact the enhanced resistance in RG-pic3 older leaves, we conducted a gene ontology (GO) analysis. GO terms associated with 69 biological processes, 6 cellular components and 54 molecular functions were significantly enriched in the upregulated DEGs in older leaves (**Supplemental Table S1A**; FDR < 0.05). GO terms in the biological processes and molecular function categories with the lowest FDR were related to defense responses and protein phosphorylation, respectively (**Figure 4C**), supporting the idea that Pic3 might regulate immunity by catalyzing dephosphorylation reactions. GO terms in the cellular component category with lowest FDR were related to subcellular localization that is consistent with our observation of Pic3 at the cell periphery (**Figure 4C**). In contrast, many of the enriched GO terms in downregulated DEGs in older leaves were related to photosynthesis (**Figure 4D** and **Supplemental Table S1B**). These results are not unexpected as photosynthesis and growth are known to be downregulated when plants allocate more resource for defense signaling (Bolton, 2009; Su et al., 2018).

We next sought to determine if certain individual upregulated DEGs significantly contribute to the enhanced resistance in older leaves of RG-pic3. We narrowed our focus for such genes to the 105 most highly upregulated DEGs in RG-pic3 older leaves (fold change ≥10, FDR < 0.05). Among these 105 genes, a cluster of genes (*Solyc12g100240*, *Solyc12g100250*, *Solyc12g100270* and *Solyc12g049030*) involved in defense-related fatty acid biosynthesis (Jeon et al., 2020) and several other potential defense related genes were selected for functional analysis (**Supplemental Table S2**). We used the agromonas approach to test the possible contribution of 11 of these genes to defense by overexpressing them in *N. benthamiana* and measuring the response to DC30000*ΔhopQ1-1ΔfliC.* None of the genes altered *N. benthamiana* resistance to DC30000*ΔhopQ1-1ΔfliC* (**Supplemental Figure S9**), suggesting the enhanced resistance in RG-pic3 older leaves is likely due to the combined effect of multiple genes/defense responses.

We found that the transcript abundance of *Bti9*, which encodes a lysin motif (LysM) receptor-like kinase (AvrPto<u>B t</u>omato-interacting <u>9</u>), is increased in older leaves of the RG-pic3 mutants compared to wild-type plants after DC3000ΔΔΔ inoculation (**Figure 4B**). The Bti9 protein has the highest sequence identity among tomato LysM-RLKs with the Arabidopsis chitin co-receptor CERK1 and is required for resistance to *P*. *syringae* and *Botrytis cinerea* (Zeng et al., 2012; Jaiswal et al., 2022). The closest Pic3 homolog protein in Arabidopsis is CIPP1, which is known to negatively regulate immunity by dephosphorylating CERK1 (Liu et al., 2018). To test if Pic3 might regulate *Pst* resistance through Bti9, we crossed a loss-of-function RG-bti9 mutant with the RG-pic3-1 mutant and generated an RG-pic3/bti9 double mutant. Upon inoculation with DC3000ΔΔΔ, the RG-bti9-1 mutant had more severe disease symptoms and slightly higher bacterial growth in older leaves than wild-type plants although the difference in bacterial populations in some experiments was not statistically significant (**Supplemental Figures S10A, S10B**, **S10C** and **S10D**). Importantly, the loss-of-function mutation in *Bti9* did not significantly compromise the increased resistance in lower leaves to DC3000ΔΔΔ in the RG-pic3/RG-bti9 plants (**Supplemental Figures S10A, S10B, S10C** and **S10D**). In addition, co-expression of Bti9 and Pic3 in leaves of *N. benthamiana* did not show any evidence of Pic3 affecting Bti9 phosphorylation during defense activation (**Supplemental Figure S11**). We therefore conclude that Bti9 does not play a detectable role in Pic3-modulated immunity to *Pst*.

### Another PP2C phosphatase, Pic12, is closely related to Pic3 and also negatively regulates the tomato immune response

In a review of our earlier RNA-Seq data (Rosli et al., 2013; Pombo et al., 2014), we noticed that the *Pic3* gene has a remarkably similar pattern of expression during the tomato PTI and ETI responses as another PP2C gene, *Pic12* (Solyc10g047290; **Supplemental Table S3**). This was also the case for their expression in response to the flagellin-deficient strain DC3000ΔΔΔ in both older and younger leaves (**Supplemental Figures S12A** and **S12B).** Interestingly, the Pic3 and Pic12 proteins share 84% amino acid identity and, are closely related to Arabidopsis CIPP1 and a defense-related PP2C (Os04g37904) in rice (Wang et al., 2012) (**Supplemental Figures S12C** and **S12D**). Together, these observations raised the possibility that Pic3 and Pic12 might have overlapping defense functions in tomato.

To investigate the role of Pic12 in the tomato defense response, we used the CRISPR/Cas9 approach described above, to generate two independent RG-pic12 homozygous mutant lines (**Figure 5A**). Both lines have mutations in the second exon of the *Pic12* gene with one (RG-pic12-1) having a 14 bp deletion and the other (RG-pic12-2) a 1 bp insertion. Each mutation causes a shift in the open reading frame resulting in predicted truncated proteins lacking the PP2C domain (**Figure 5A** and **Supplemental Figure S13A).** The RG-pic12 mutant plants had wild-type morphology at 4 weeks old (**Figure 5A**). At 8 weeks old, the RG-pic12 plants were slightly shorter than wild-type and at 12 weeks old the plants showed earlier senescence than wild-type (**Supplemental Figures S13B** and **S13C**).

**Figure 5.**
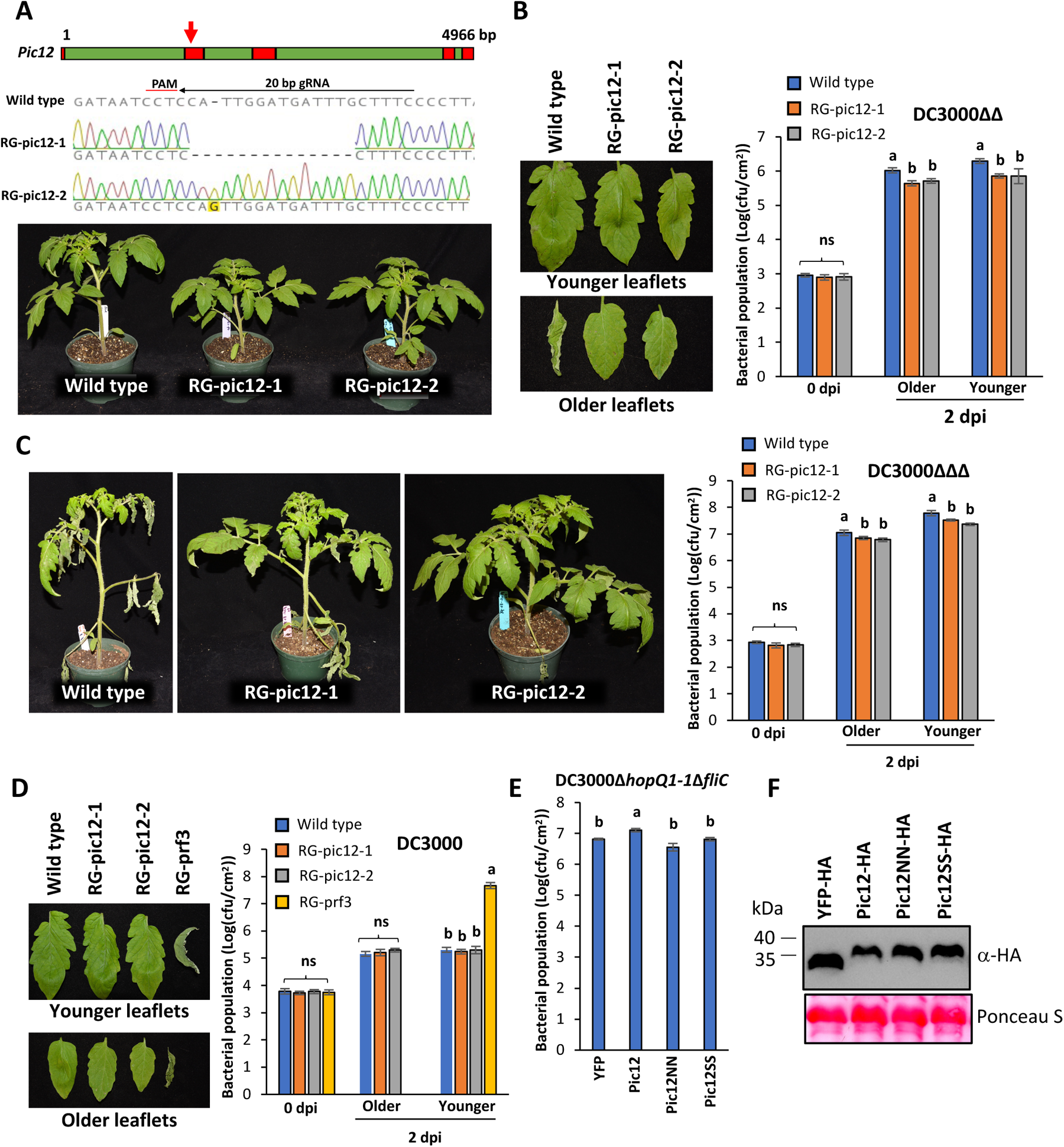
Tomato loss-of-function RG-pic12 mutants show increased disease resistance. **A)** Top panel, schematic showing gene structure of *Pic12* (Solyc10g047290) including exons (red) and introns (green), arrow points to CRISPR/Cas9 gRNA target site; Middle panel, sequencing chromatogram of two missense mutant alleles generated by CRISPR/Cas9; Bottom panel, images of 4-week-old wild-type (Rio Grande-PtoR; RG-PtoR) and RG-pic12 mutant plants. **B)** Wild-type and RG-pic12 plants were vacuum-infiltrated with 5×10^4^ cfu/mL DC3000Δ*avrPto*Δ*avrPtoB* (DC3000ΔΔ), photographed at 4 days post-inoculation (dpi). **C)** Wild-type and RG-pic12 plants were vacuum-infiltrated with 5×10^4^ cfu/mL DC3000Δ*avrPto*Δ*avrPtoB*Δ*fliC* (DC3000ΔΔΔ), and photographed at 4 dpi. **D)** Wild-type, RG-pic12 and RG-prf3 plants were vacuum-infiltrated with 5×10^5^ cfu/mL DC3000, and photographed at 4 dpi. ‘Older’ refers to 1^st^ or 2^nd^ true leaf and ‘younger’ refers to 4^th^ or 5^th^ true leaf. Bacterial populations were measured at 0 and 2 dpi. **E)** Leaves of 5-week-old *N. benthamiana* plants were syringe-infiltrated with Agrobacterium (1D1249) strains (OD_600_ = 0.6) carrying a binary expression vector (pGWB414) expressing 3× HA-tagged wild-type Pic12, a phosphatase-inactive variant Pic12NN (D69N and D231N substitutions), or a Pic12SS variant with substitutions in the palmitoylation motif (C5S and C6S substitutions) or a YFP control. Two days later, the same agroinfiltrated spots were syringe-infiltrated with 5×10^4^ cfu/mL DC3000*ΔhopQ1-1ΔfliC.* DC3000*ΔhopQ1-1ΔfliC* populations were measured at 2 days after the second infiltration. **F)** Western blotting analysis confirmed that proteins in (E) were expressed. Bars show means ± standard deviation, n = 3 or 4. cfu, colony-forming units. Different letters indicate significant differences based on a one-way ANOVA followed by Tukey’s Honest Significant Difference post hoc test (P < 0.05). ANOVA was performed separately for each time point and leaf type. ns, no significant difference. Experiments (B, C and D) were repeated three times with similar results. Experiment (E) was repeated twice with similar results.

We tested defense-related responses of the RG-pic12 plants by inoculating them and wild-type plants with the three *Pst* DC3000 strains described earlier. Unlike the RG-pic3 mutant lines, the two RG-pic12 lines showed increased resistance to DC3000ΔΔ and DC3000ΔΔΔ in both older and younger leaves, as shown by less disease symptoms and lower bacterial populations (**Figures 5B** and **5C**). Disease symptoms and bacterial populations in RG-pic12 and wild-type plants after inoculation with DC3000 were similar (**Figure 5D**). Together, these observations suggest that Pic12, like Pic3, regulates flagellin-independent defense responses but does not appear to affect Pto/Prf-mediated ETI.

The Pic12 protein has two cysteines (C5 and C6) that are predicted to be palmitoylated and two conserved aspartic acids (D69 and D231) expected to be required for its phosphatase activity (**Supplemental Figure S14A**). To test the importance of these residues, we made serine substitutions for the two cysteines (C5S and C6S; Pic12SS) and asparagine substitutions for the two aspartic acid residues (D69N and D231N; Pic12NN). The wild-type Pic12 and each variant were transiently expressed in leaves of *N. benthamiana* and the effect on subsequent inoculation with DC3000*ΔhopQ1-1ΔfliC* was measured. Expression of wild-type Pic12 but not Pic12SS or Pic12NN increased *N. benthamiana* susceptibility to DC3000*ΔhopQ1-1ΔfliC* relative to a YFP control (**Figure 5E**). An immunoblot showed that all proteins were expressed at a similar level (**Figure 5F**). These data indicate that like Pic3, the palmitoylation motif and phosphatase activity are required for Pic12-mediated negative regulation of plant defense.

### Pic12 is localized to the cell periphery and does not affect flagellin-induced MAPK activation or ROS production

Because the substitutions in the Pic12 palmitoylation motif compromised its defense function, we hypothesized that, like with Pic3, this motif is required for Pic12 subcellular localization. We fused wild-type Pic12 and Pic12SS to a C-terminus YFP tag and visualized their localization in the plant cell by transiently expressing each protein (along with a YFP control) in leaves of *N. benthamiana* (**Supplemental Figure S14B**). Like Pic3, wild-type Pic12-YFP protein was primarily localized to cell periphery. However, the Pic12-YFP fluorescence at the periphery largely disappeared in leaves expressing Pic12SS-YFP (**Supplemental Figure S14B**), suggesting the palmitoylation motif is required for proper localization of Pic12. An immunoblot showed that all proteins were expressed in these experiments, though Pic12-YFP was less abundant (**Supplemental Figure S14C**). As with Pic3, Pic12 localization was not changed after DC3000*ΔhopQ1-1* inoculation or flg22 treatment (**Supplemental Figure S14D**).

To test if Pic12 plays a role in flagellin-mediated PTI, we measured MAPK activation and ROS production in older and younger leaves of RG-pic12 and wild-type plants after either flg22 or flgII-28 treatment. We observed similar levels of MAPK activation and ROS production in both RG-pic12 and wild-type plants in these experiments (**Supplemental Figures S15A**, **S15B**, and **S15C**). Pic12 therefore does not appear to regulate two host responses associated with flagellin-mediated PTI.

### Pic3 and Pic12 negatively regulate resistance to another bacterial pathogen, *Xanthomonas euvesicatoria*

Our experiments to this point had focused on the response of the RG-pic3 and RG-pic12 mutants to different strains of DC3000 and supported a role of both PP2Cs in the negative regulation of plant defenses against this bacterial pathogen. To extend these observations beyond *Pst*, we inoculated RG-pic3, RG-pic12 and wild-type plants with another bacterial foliar pathogen, *Xanthomonas euvesicatoria* (*Xe*), the causative agent of bacterial spot disease of tomato and measured bacterial populations in older and younger leaves. Like the responses to *Pst*, RG-pic3 plants showed enhanced resistance in older leaves to *Xe*, while RG-pic12 plants showed enhanced resistance to this pathogen in both older and younger leaves (**Supplemental Figure S16**).

### Pic3 and Pic12 additively contribute to bacterial disease resistance in tomato

Except for the RG-pic3 mutants showing enhanced resistance only in older leaves compared to the enhanced resistance in both older and younger leaves of RG-pic12 plants, similar phenotypes of the mutant lines, the relatedness of the Pic3 and Pic12 proteins, and their similar gene expression patterns suggested that the two PP2Cs might have at least partially redundant/additive functions in negatively regulating flagellin-independent immunity. To test this possibility, we crossed RG-pic3-1 and RG-pic12-2 to generate an RG-pic3/RG-pic12 double mutant. The double mutant began to show altered morphology at 4 weeks-old with noticeably deformed older leaves (**Figure 6A**). At 8 weeks-old, leaves of RG-pic3/RG-pic12 plants showed pronounced early senescence compared to wild-type and single RG-pic3 or RG-pic12 plants (**Figure 6A**) and they produced very few viable seeds. Upon inoculation with DC3000ΔΔΔ, RG-pic3/RG-pic12 plants supported smaller bacterial populations in older leaves than the RG-pic3 or RG-pic12 single mutants, although disease symptoms in the RG-pic12 plants were as severe as in wild-type plants with this DC3000 strain (**Figures 6B** and **6C**). In younger leaves the RG-pic3/RG-pic12 and RG-pic12 plants both supported lower bacterial populations and showed less severe disease symptoms (**Figures 6B** and **6C**). Collectively, with respect to bacterial growth, these observations indicate that in older leaves Pic3 and Pic12 act in a partially redundant manner, whereas in younger leaves Pic12 acts without Pic3. MAPK activation after flg22 or flgII-28 treatment in RG-pic3/RG-pic12 plants was not different from wild-type or RG-pic3 or RG-pic12 single mutant plants in older and younger leaves (**Supplemental Figure S17**), further indicating that neither Pic3 nor Pic12 affect this flagellin-associated PTI response.

**Figure 6.**
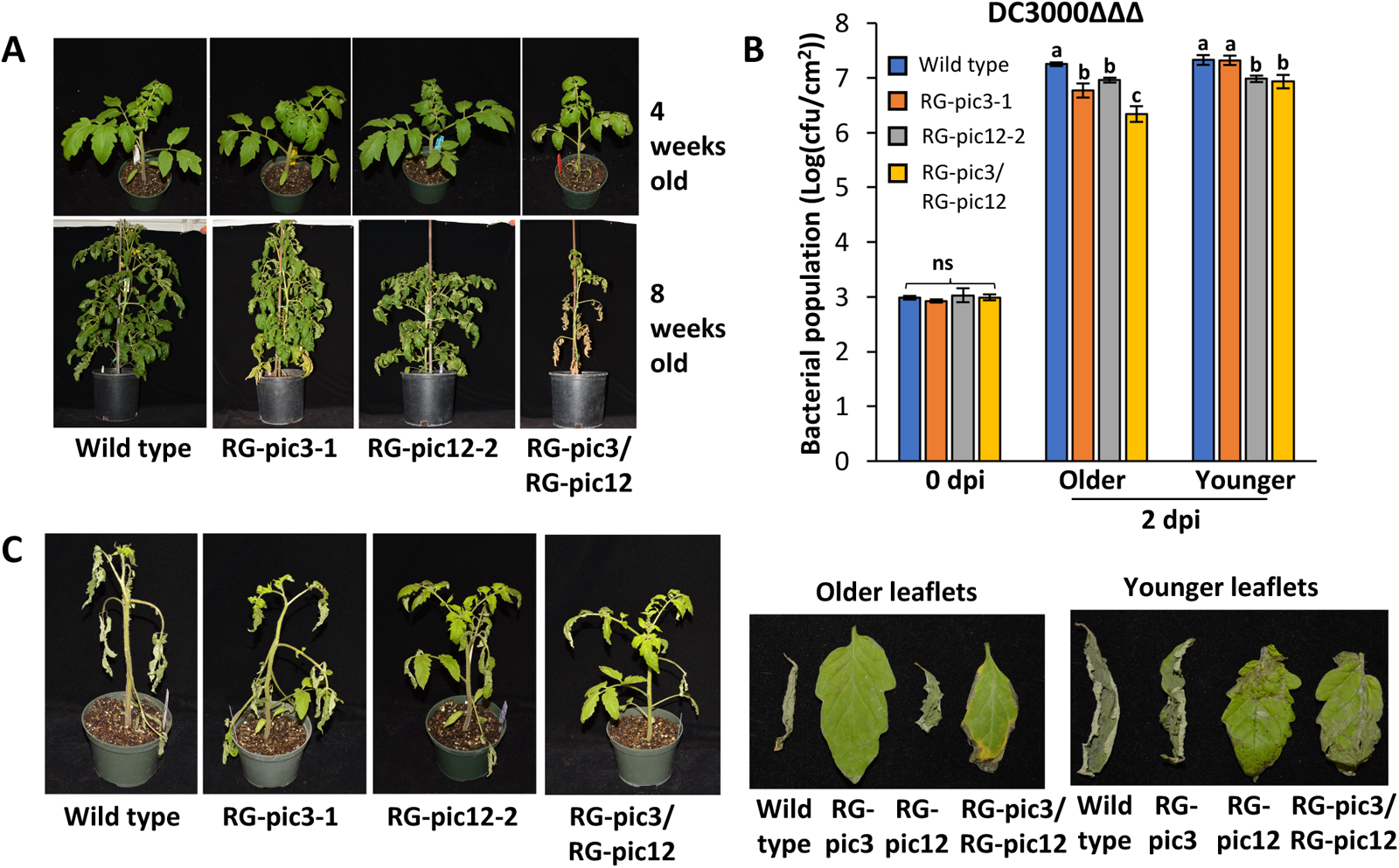
Pic3 and Pic12 additively contribute to disease resistance to *Pst*. **A)** Morphology of 4-and 8-week-old wild-type, RG-pic3-1, RG-pic12-2 and RG-pic3/RG-pic12 mutant plants. **B)** Wild-type, RG-pic3-1, RG-pic12-2 and RG-pic3/RG-pic12 plants were vacuum-infiltrated with 5×10^4^ cfu/mL DC3000Δ*avrPto*Δ*avrPtoB*Δ*fliC* (DC3000ΔΔΔ). Bacterial populations were measured at 0 and 2 days post-inoculation (dpi). ‘Older’ refers to 1^st^ or 2^nd^ true leaf and ‘younger’ refers to 4^th^ or 5^th^ true leaf. Bars show means ± standard deviation, n = 4. cfu, colony-forming units. Different letters indicate significant differences based on a one-way ANOVA followed by Tukey’s Honest Significant Difference post hoc test (P < 0.05). ANOVA was performed separately for each time point and leaf type. ns, no significant difference. **C)** Disease symptoms at 4 dpi on whole plants or the older and younger leaves of indicated genotypes related to experiment in (B). Inoculation experiments were repeated three times with similar results.

## Discussion

Protein phosphorylation is a reversable posttranslational modification that can regulate protein conformation, stability, activity, and localization. In many biological processes, protein kinases and phosphatases act as a molecular switch by turning protein phosphorylation ‘on’ and ‘off’ (Zhang et al., 2023). In defense responses, plants rely on transmembrane cell-surface and intracellular receptors to sense the presence of pathogens. Many of the surface receptors have an intracellular protein kinase domain and they transduce outside stimuli into coordinated cellular responses by phosphorylating specific substrates (Bi et al., 2018; DeFalco and Zipfel, 2021). As the environment changes, plants adjust their responses by modulating cellular protein phosphorylation status via protein phosphatases (Erickson et al., 2022; Sobol et al., 2022). Similarly, some intracellular NLR receptors rely on protein kinases to activate robust immune responses upon recognition of specific pathogen ‘effector’ proteins (del Pozo, Pedley and Martin, 2004; Pedley and Martin, 2004; Roberts et al., 2019; Zhang and Zhang, 2022). While much progress has been made in understanding the role of protein kinases in plant immunity, less is known about how protein phosphatases regulate defense signaling. PP2C protein phosphatases constitute a large family in plants and a few studies have shown them to play a role in plant immunity (Schweighofer et al., 2007; Park et al., 2008; Couto et al., 2016; Giska and Martin, 2019). In this study, we used CRISPR/Cas9 gene editing technology in tomato to generate loss-of-function mutations in a collection of <u>P</u>P2C-immunity associated <u>c</u>andidate (Pic) genes and investigated their possible roles in bacterial resistance. We discovered and characterized two PP2Cs that negatively regulate the plant immune response in a leaf age-specific manner.

Our screening of the Pic mutants for their response to three *Pst* DC3000 strains led us to discover two PP2C phosphatases, Pic3 and Pic12, that negatively regulate disease resistance to two different bacterial pathogens. Interestingly, the enhanced resistance in RG-pic3 (but not RG-pic12) plants is observed only in older leaves and not in younger leaves. Consistent with this observation, our RNA-seq analysis of RG-pic3 vs. wild-type plants revealed significantly more DEGs in older than in younger leaves. This difference in older and younger leaves is reminiscent of the leaf age-dependent response to *Pst* DC3000 seen in Arabidopsis *fls2* mutants (Zipfel et al., 2004). In that case, the *fls2* mutant plants show increased disease susceptibility in younger leaves as compared to older leaves (Zipfel et al., 2004). However, the different disease symptoms seen in older and younger leaves of the RG-pic3 lines is not likely to involve Fls2 (or Fls3) because RG-pic3 plants also show enhanced resistance in older leaves to DC3000ΔΔΔ, which does not produce flagellin. In addition, MAPK activation and ROS production were mostly similar in RG-pic3, RG-pic12 and wild-type plants after various PAMP treatments, further indicating that the two phosphatases do not have a significant effect on flagellin-activated defense pathways.

The localization of Pic3 and Pic12 at the cell periphery suggests that they might regulate defense responses by dephosphorylating immunity-associated proteins near plasma membrane. The GO term analysis showed clear enrichment for terms such as “membrane”, “phosphorylation”, and “defense” in the upregulated DEGs in older leaves of RG-pic3 vs. wild-type plants, further supporting this hypothesis. We used the agromonas assay to investigate the role of some of these upregulated DEGs in the Pic3-mediated response. However, overexpression of 11 individual candidate genes (selected from 105 DEGs with highest fold change) did not alter resistance to DC3000 in leaves of *N. benthamiana*. Although we do see an effect on DC3000 when Pic3 and Pic12 are used in the agromonas assay, it seems likely that the enhanced defense seen in older leaves of RG-pic3 plants is due to the upregulation of an array of genes working synergistically or additively. Our RNA-seq data did show that several known PTI-associated genes are among the upregulated DEGs in older RG-pic3 leaves, and one or more of these genes could conceivably contribute to the enhanced resistance we observed.

Pic3 and Pic12 share similarities in amino acid sequence, gene expression pattern, localization, and both negatively regulate defense in a flagellin independent manner. However, there is evidence that Pic3 has a slightly different regulatory role than Pic12. First, RG-pic3 mutants show increased resistance only in older leaves whereas RG-pic12 mutants exhibit enhanced resistance in both older and younger leaves. Second, RG-pic3, but not RG-pic12, mutants show slightly enhanced resistance to DC3000. Third, the RG-pic3, but not RG-pic12, plants develop early senescence when they are two-months old. Fourth, flgII-28 induced ROS is higher in older leaves of RG-pic3 than in wild-type plants, while Pic12 does not have an effect on flgII-28 induced ROS. Fifth, the RG-pic3/RG-pic12 plants show a stronger resistance to DC3000ΔΔΔ than the RG-pic3 or RG-pic12 single mutants. Interestingly, the pronounced spontaneous cell death seen in unchallenged RG-pic3/RG-pic12 plants is not seen in RG-pic3 or RG-pic12 single mutants. Collectively, these observations suggest that Pic3 and Pic12 act in at least a partially redundant fashion and conceivably target some common and some different substrates to regulate defense responses.

Phylogenetic analysis of Pic3, Pic12 and related PP2C phosphatases revealed that Pic3 and Pic12 (along with Solyc04g074190, but this gene was not induced after PAMP treatments) are closely related to *Arabidopsis* phosphatases AtCIPP1 and AtPAPP2C, as well as rice Os04g37904 (**Supplemental Figure S12C**). CIPP1 has been shown to negatively regulate defense by dephosphorylating CERK1, and both AtPAPP2C and Os04g37904 have been implicated in regulating salicylic acid (SA)-dependent immunity (Wang et al., 2012; Liu et al., 2018). Our previous work and a recent study by others have demonstrated that the LysM-RLK Bti9 (also named LYK1) is the closest homology of CERK1 in tomato (Zeng et al., 2012; Jaiswal et al., 2022). Bti9 RNAi-silenced plants show increased susceptibility to *Pst* specifically in older leaves (Zeng et al., 2012). However, in our present epistatic analysis of an RG-pic3/RG-bti9 double mutant we found that the absence of Bti9 had little effect on the enhanced resistance seen in older leaves of RG-pic3 (**Supplemental Figure S10**). Moreover, we found that co-expression of Pic3 and Bti9 in *N. benthamiana* leaves does not affect Bti9 phosphorylation status in plant cells (**Supplemental Figure S11**). These observations suggest that Pic3 does not play a detectable role in Bti9-associated defense. Investigation of other possible substrates of Pic3 and Pic12 will be a focus of our future work.

We considered the possible role of salicylic acid (SA) and abscisic acid (ABA) in Pic3-associated defense. Isochorismate synthase, a key enzyme in SA biosynthesis, is encoded by the *ICS* gene which, in *Arabidopsis*, is transcriptionally upregulated during many defense responses (Macaulay et al., 2017). However, the transcript abundance of the single *ICS* gene that is present in tomato (Changwal et al., 2021) was not significantly increased in RG-pic3 plants compared to wild-type plants after *Pst* inoculation based on our RNA-seq data. Regarding ABA, no members of the PP2C clade D PP2Cs, to which Pic3 and Pic12 belong, have yet been implicated in ABA signaling (Qiu et al., 2022). Additionally, we observed no effect on the expression of two ABA reporter genes in the RG-pic3 mutant plants. (**Supplemental Figure S5**), suggesting ABA is not involved in the enhanced resistance seen in RG-pic3 plants. Further research is likely needed to understand if Pic3 or Pic12 contributes to SA, ABA or other immunity-associated hormone pathways during *Pst* attack.

Based on our current data, we propose that Pic3 and Pic12 act at the cell periphery to dephosphorylate common as well as some different pattern recognition receptors (PRRs) or their downstream signaling component(s) to regulate immunity. Several of our observations support this hypothesis. First, the palmitoylation motifs of Pic3 and Pic12 are required for localization to the cell periphery and disruption of palmitoylation motifs abolished the ability of Pic3 and Pic12 proteins to promote *Pst* growth in *N. benthamiana* plants, suggesting they both act at or near the plasma membrane. Second, loss of either Pic3 or Pic12 leads to increased resistance to *Pst*, and overexpression of either Pic3 or Pic12 (but not phosphatase-inactive variants) renders *N. benthamiana* plants more susceptible to *Pst*. These responses are independent of flagellin recognition, suggesting that Pic3 and Pic12 might regulate immunity triggered by other PAMPs. Pic3 and Pic12 could also act on certain conserved proteins downstream of PRRs. Because RG-pic3 and RG-pic12 mutants have some different defense phenotypes, it is possible that they target some unique substrates. Identifying the substrates of Pic3 and Pic12 is an important next step for further understanding how Pic3 and Pic12 regulate immunity at the molecular and protein levels. Overall, our research has revealed a key role for two PP2C phosphatases in plant immunity and sets the stage for future study of the substrates they target to negatively regulate immune responses.

## Materials and Methods

### Generation of RG-pic mutant lines

For each *Pic* gene, we designed one or two guide RNAs (gRNAs) targeting the exons near the 5’ terminus using the software Geneious R11 (Kearse et al., 2012). Each gRNA cassette was cloned into a Cas9-expressing binary vector (p201N:Cas9) by Gibson assembly as described previously (Jacobs and Martin, 2016). The p201N:Cas9 construct carrying each gRNA was introduced into *Agrobacterium tumefaciens* strain AGL1 through electroporation. For *Pic* genes that were targeted by two gRNAs, Agrobacterium cells containing each gRNA/Cas9 construct were pooled together and used for transformation into the tomato cultivar Rio Grande-PtoR (RG-PtoR, which has the *Pto* and *Prf* genes). Tomato transformation was carried out at the Center for Plant Biotechnology Research at the Boyce Thompson Institute for Plant Research. Transgenic plants were confirmed by PCR using specific primers (**Supplemental Table S4**) and sequencing at the Biotechnology Resource Center at Cornell University. Geneious R11 and the web-based tool TIDE (Tracking of Indels by Decomposition, https://tide.deskgen.com; (Brinkman et al., 2014)) were used to determine the mutation type and frequency using the sequencing files (ab1 format).

### Off-target Mutation Detection

For Pic3 and Pic12 loss-of-function mutant lines, two potential off-target sites (with the highest probabilities) predicted by CRISPR-P v2 (Liu et al., 2017) were tested. Five plants from RG-pic3-1 or RG-pic12-1 T3 populations were randomly selected and PCR was conducted to amplify the DNA fragments that flank these predicted off-target sites using specific primers (**Supplemental Table S4**). PCR products were sequenced (Sanger sequencing) and compared with reference gene sequences from the tomato genome database (https://solgenomics.net). No alterations of any of the potential off-targets was observed which aligns with our characterization of off-targets in a larger collection of tomato mutants generated by CRISPR/Cas9 (Zhang et al., 2020).

### Disease Inoculation Assay

Four-to five-week-old tomato plants (a stage showing no senescence in RG-pic3 or RG-pic12) were used for disease assays. Tomato plants were submerged into a suspension of *P. syringae* pv. *tomato* strains DC3000, DC3000Δ*avrPto*Δ*avrPtoB* (DC3000ΔΔ), DC3000Δ*avrPto*Δ*avrPtoB*Δ*fliC* (DC3000ΔΔΔ), or *Xanthomonas euvesicatoria* strain 85-10 and the whole plant was uniformly inoculated using vacuum infiltration. Inoculation titers ranged from 2x10^4^ to 1x10^7^ cfu/mL and are indicated in the figure legends. Three or four plants per line were tested with each bacterial strain. Bacterial populations were measured at 3 hr (for 0 days post-inoculation; dpi) and 2 dpi. Disease symptoms were photographed at 3 to 8 days after bacterial infection.

### Construct Generation

The coding region of each gene was amplified from tomato cDNA using KOD hot start DNA polymerase (Thermo Fisher Scientific) and gene-specific primers (**Supplemental Table S4**) and first cloned into pENTR/D-TOPO entry vector following manufacturer’s instructions (Thermo Fisher Scientific). The gene expression cassette in pENTR/D-TOPO was then cloned into the destination vector pGWB414, pGWB417 or pGWB541 (Nakagawa et al., 2007) via recombination reactions using LR Clonase II (Thermo Fisher Scientific). Vectors were confirmed by Sanger sequencing and transformed into *Agrobacterium tumefaciens* strain 1D1249, GV2260 or GV3101 for transient expression in *N. benthamiana*.

Palmitoylation motifs were predicted using the online tool GPS-Palm (https://gpspalm.biocuckoo.cn/; (Ning et al., 2021)). Nucleotide changes leading to amino acid substitutions for Pic3 and Pic12 were first created in pENTR/D-TOPO (which contained wild-type Pic3 or Pic12) vectors by using the Q5 site-directed mutagenesis kit (New England Biolabs) with specific primers (**Supplemental Table S4**). The resulting mutated vectors were then cloned into the destination vector pGWB414, or pGWB541 through an LR reaction.

### Agromonas Assay

The agromonas assays were performed as described previously (Buscaill et al., 2021). Briefly, Agrobacterium strain 1D1249 carrying a binary vector (pGWB414 or pGWB417) expressing the gene of interest were syringe infiltrated into leaves of 5-week-old *N. benthamiana* plants. Two days later, the same agroinfiltrated spots were syringe infiltrated with either *Pst* DC3000Δ*hopQ1-1* or DC3000Δ*hopQ1-1ΔfliC* at 5x10^4^ cfu/mL. Bacterial populations were measured by serial dilutions on LB medium supplemented with 10 µg/mL cetrimide, 10 µg/mL fucidin, and 50 µg/mL cephaloridine (CFC; Oxoid C-F-C Supplement) at 2 days after *Pst* inoculation.

### Confocal Microscopy

*N. benthamiana* leaves were agroinfiltrated to transiently express the protein of interest and examined with a Leica TCS SP5 Laser Scanning Confocal Microscope (Leica Microsystems, Exton, PA, USA) using a 63x water immersion objective NA 1.2 (HCX PL APO). YFP was excited using a 488-nm laser line from an argon laser line and collected between 525 nm and 550 nm using an HyD detector. Images were processed using Leica LAS-AF software version 2.7.3 (Leica Microsystems). Microscopy analysis was conducted at Plant Cell Imaging Center at the Boyce Thompson Institute for Plant Research.

### MAPK Phosphorylation Assay

MAPK assays were performed as described previously (Zhang et al., 2020). Briefly, six leaf discs from 4-week-old plants were floated in water overnight to let the wound response subside. The leaf discs were then incubated in 10 nM of flg22, 25 nM of flgII-28, 50 nM of csp22 or water (negative control) for 10 min, and flash frozen in liquid nitrogen. Protein was extracted using a buffer containing 50mM of Tris-HCl at pH 7.5, 10% (v/v) glycerol, 2mM of EDTA, 1% (v/v) Triton X-100, 5 mM of DTT (dithiothreitol), 1% (v/v) protease inhibitor cocktail (Sigma-Aldrich), and 0.5% (v/v) phosphatase inhibitor cocktail 2 (Sigma-Aldrich). MAPK phosphorylation was determined using an antiphospho-p44/42 MAPK (Erk1/2) antibody (anti-pMAPK; Cell Signaling).

### Immunoblotting

Total protein was extracted from *N. benthamiana* leaves using protein extraction buffer containing 50 mM Tris-HCl (pH 7.5), 10% glycerol, 150 mM NaCl, 10 mM MgCl_2_, 5 mM EDTA, 5 mM DTT (dithiothreitol), and 1% (v/v) protease inhibitor cocktail (Sigma-Aldrich). A 12 µL soluble protein solution mixed with 4 µL Laemmli sample buffer was boiled at 95°C for 6 min before loading for gel electrophoresis. Protein was separated by 8%–12% sodium dodecyl sulfate–polyacrylamide gel electrophoresis (SDS-PAGE) and blotted on polyvinylidene difluoride membrane (Merck Millipore). The membranes were exposed to anti-HA (1:5000; v/v), anti-MYC (1:10000; v/v), or anti-GFP (1:5000; v/v) primary antibody and corresponding anti-rat-HRP (1:10000; v/v; for HA) or anti-mouse-HRP (1:10000; v/v; for MYC and GFP) secondary antibody and developed with Pierce ECL plus substrate (Thermo Fisher Scientific) for 5 min. Images were acquired by using Bio-Rad ChemiDoc Imaging system (Bio-Rad).

### Phos-tag phosphoprotein separation

Phosphoprotein separation was carried out using Phos-tag SDS PAGE as described previously (Liu et al., 2023) with slight modification. Briefly, total protein was extracted from *N. benthamiana* leaves (co-expressing Bti9-MYC with Pic3-HA or YFP-HA proteins) using protein extraction buffer containing 50 mM HEPES (4-(2-hydroxyethyl)-1-piperazineethanesulfonic acid pH 7.5), 10% glycerol, 150 mM NaCl, 50 mM β-glycerophosphate, 2 mM DTT (dithiothreitol), 1% (v/v) Triton X-100, 1% (v/v) protease inhibitor cocktail (Sigma-Aldrich) and 0.5% (v/v) phosphatase inhibitor cocktail 2 (Sigma-Aldrich). Protein extracts were separated in an 8% SDS-PAGE gel containing 40 µM Phos-tag (Fujifilm Wako Chemicals), 150 µM MnCl_2_ and 0.38 M Tris-HCl (pH 8.4). Note that the protein ladder is incompatible with Phos-tag gel and was not used in these experiments. The proteins were blotted on polyvinylidene difluoride membrane (Merck Millipore) and Bti9-MYC was detected with an anti-MYC antibody.

### ROS Assay

ROS production was measured as described previously (Hind et al., 2016). Briefly, leaf discs were collected and floated in 200 µL water in a 96-well Nunc flat-bottom microplate overnight (16 h). Water was then removed and replaced with a 100 µL solution containing 100 nM of flg22 (QRLSTGSRINSAKDDAAGLQIA) or 100 nM of flgII-28 (ESTNILQRMRELAVQSRNDSNSSTDRDA), in combination with 34 mg/mL of luminol (Sigma-Aldrich) and 20 mg/mL of horseradish peroxidase. ROS production was monitored over 45 min using a Synergy-2 microplate reader (BioTek). Three plants per line and two to four discs per plant were tested in each experiment.

### qRT-PCR

Wild-type 4-week-old RG-PtoR plants were vacuum infiltrated with 5x10^6^ cfu/mL DC3000Δ*avrPto*Δ*avrPtoB*Δ*fliC* or 10 mM MgCl_2_. At least three plants were used for each treatment. After 6 h, around 50 mg of tissue from the older (1^st^ or 2^nd^) and younger (4^th^ or 5^th^) leaves from each plant were collected and flash frozen in liquid N_2_. Total RNA was isolated using an RNeasy Plant Mini Kit (Qiagen). Total RNA (about 3 µg) was treated with TURBO DNA-free DNase (Thermo Fisher Scientific) for 45 min at 37°C. First-strand complementary DNA was then synthesized using the above RNA via SuperScript III and Oligo(dT)20 primer (Thermo Fisher Scientific). Quantitative PCR was performed with specific primers (**Supplemental Table S4**) using the QuantStudio 6 Flex Real-Time PCR System (Thermo Fisher Scientific) and cycling conditions for PCR were 50°C for 2 min, 95°C for 10 min, and followed by 40 cycles of 95°C for 30 s, 60°C for 1 min. Tomato *SlArd2* (*Solyc01g104170*) was used as the reference gene (Pombo et al., 2017).

### RNA-Seq

Four-week-old wild-type RG-PtoR and RG-pic3-1, RG-pic3-2 mutant plants were vacuum infiltrated with 5x10^6^ cfu/mL DC3000Δ*avrPto*Δ*avrPtoB*Δ*fliC*. Six leaf discs (about 50 mg) from older (1^st^ or 2^nd^) or younger (4^th^ and 5^th^) leaves were collected at 6 h after infiltration. Total RNA was isolated using an RNeasy Plant Mini Kit (Qiagen) and then was treated with TURBO DNA-free DNase (Thermo Fisher Scientific) for 45 min at 37°C. RNA libraries were prepared and sequenced on an Illumina HiSeq 4000 system. Raw RNA-seq reads were processed to remove adaptors and low-quality sequences using Trimmomatic (version 0.39) with default parameters (Bolger, Lohse and Usadel, 2014). The remaining cleaned reads were aligned to the ribosomal RNA database (Quast et al., 2013) using bowtie (version 1.1.2) (Langmead, 2010) allowing up to three mismatches, and those aligned were discarded. The remaining cleaned reads were mapped to the tomato reference genome (SL4.0 and ITAG4.1) using HISAT2 (version 2.1.0) (Kim et al., 2019) with default parameters. Based on the alignments, raw read counts for each gene were calculated using HTSeq-count (Anders, Pyl and Huber, 2015) and normalized to FPKM. Raw read counts were then fed to DESeq2 (Love, Huber and Anders, 2014) to identify differentially expressed genes, with a cutoff of adjusted P (FDR) < 0.05 and fold change ≥2. Raw RNA-Seq reads have been deposited in the NCBI BioProject database under the accession number PRJNA1079816.

## Acknowledgments

We thank Tara McCrudden, Julie Bell, Kelly Jackson, Philip Ricci and Nick Vail for plant care, Joyce Van Eck, Qingzhen Jiang and Tish Keen for tomato transformation, and Sergey Ivanov and Mamta Srivastava for assistance with confocal imaging. JC thanks Professor Amir Sharon for supporting his research at Tel Aviv University. RES was funded by NSF Research for Undergraduates award DBI-1850796. This work was supported by grant IS-5362-21CR to GSe and GBM from the United States-Israel Binational Agricultural Research and Development Fund (BARD). This paper is dedicated to the memory of Guido Sessa.

## Author contributions

FX, GSe, GSo and GBM conceived and designed the experiments. FX and NZ designed gRNAs and constructed vectors. FX performed genotyping and phenotyping experiments. FX, GBM, GSe and GSo analyzed the data. RS and JC helped with phenotyping experiments. XT and ZF performed RNA-Seq data analysis. FX and GBM interpreted the data and wrote the manuscript. All the authors, except GSe, read and approved the manuscript.

