## Supplemental Data for "Related PP2C phosphatases Pic3 and Pic12 negatively regulate immunity in tomato to *Pseudomonas syringae*"

**A**

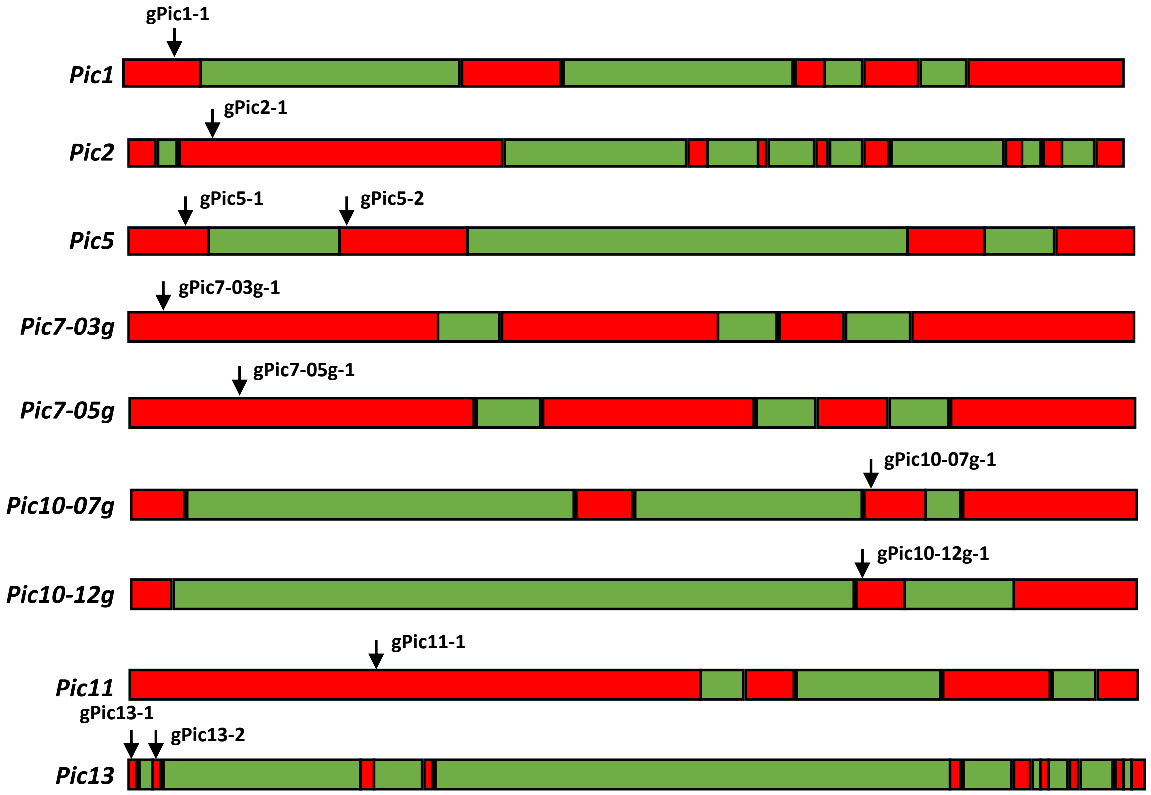


**B**

| Mutant line | Gene name | Solyc ID | Mutation | gRNA that induced the mutation |
|---|---|---|---|---|
| RG-pic1 | *Pic1* | Solyc07g066260 | +1 bp | gPic1-1 (GTGTAGGCAAGCACGGGAGG) |
| RG-pic2-1 | *Pic2* | Solyc01g105020 | -1 bp | gPic2-1 (GAAGTAATAGAATTCTCTGG) |
| RG-pic2-2 | *Pic2* | Solyc01g105020 | -11 bp | gPic2-1 (GAAGTAATAGAATTCTCTGG) |
| RG-pic5-1 | *Pic5* | Solyc02g092750 | -4 bp | gPic5-2 (GGAAGAAGGAGGATTGTCTG) |
| RG-pic5-2 | *Pic5* | Solyc02g092750 | -8 bp | gPic5-1 (GGTGTTTATGATGGTCATGG) |
| RG-pic7 | *Pic7-03g* | Solyc03g096670 | +1 bp | gPic7-03g-1 (GCTTCCGTACTCGGTTCGGT) |
| | *Pic7-05g* | Solyc05g052980 | -37 bp | gPic7-05g-1 (GTTGCAGCTGCAACATCGGT) |
| RG-pic10-1 | *Pic10-07g* | Solyc07g007220 | +1 bp | gPic10-07g-1 (CAGGTCTTTGATGGACATGG) |
| | *Pic10-12g* | Solyc12g010450 | -1 bp | gPic10-12g-1 (CAGGTCTTTGATGGACATGG) |
| RG-pic10-2 | *Pic10-07g* | Solyc07g007220 | -195 bp | gPic10-07g-1 (CAGGTCTTTGATGGACATGG) |
| | *Pic10-12g* | Solyc12g010450 | -787 bp, + 171 bp | gPic10-12g-1 (CAGGTCTTTGATGGACATGG) |
| RG-pic11 | *Pic11* | Solyc09g010780 | -75 bp | gPic11-1 (TCAAACACGGGCTCTGGATT) |
| RG-pic13-1 | *Pic13* | Solyc12g042570 | -4 bp | gPic13-1 (GCATGCAGTTCATCCAAATT) |
| RG-pic13-2 | *Pic13* | Solyc12g042570 | -2 bp | gPic13-2 (GAGAATGACAGACTAAGATA) |

**Supplemental Figure S1. Summary of the mutations in the *Pic* CRISPR lines. A)** Structure of each *Pic* gene showing exons (red) and introns (green). The arrows point to the CRISPR/Cas9 gRNA target site. **B)** RG-Pic CRISPR mutant lines and detailed gRNA, mutation information. "+ 1 bp" means 1 base pair insertion and "- 1 bp" indicates 1 base pair deletion. In the RG-pic7 mutant, two PP2C genes *Pic7-03g* and *Pic7-05g* (which share high sequence similarity) were mutated via CRISPR/Cas9 guide RNA gPic7-03g-1 and gPic7-05g-1, respectively. Similarly, in the RG-pic10-1 or RG-pic10-2 mutants, two PP2C genes *Pic10-07g* and *Pic10-12g* (which share high sequence similarity) were mutated via CRISPR/Cas9 guide RNA gPic10-07g-1 and gPic10-12g-1, respectively. In all other mutants, only one PP2C *Pic* gene was edited.

**A**

**RG-PtoR:** MDNLCCFNSKFSKLAGGRSSCSSGKGRSNQGPTKYGFSLVKGKANHPMEDYHVSKFVQL
HGHELGLFAIYDGHLGDSVPAYLQKHLFSNILNEEDFRNDPHRAILKAYERTDQAILSHSPD
LGRGGSTAVTAILINGRKLWVANVGDSRAVLSRRGQAIQLSIDHEPNTERDDIENRGGFVS
NMPGDVARVNGQLAVSRAFGDKNLKSHLSSDPDVTNADVDAETDLLILASDGLWKVMS
NQEAVDIVRKVKDPEKAAKQLAIEALVRESKDDISCIVVRFKG*

**RG-pic3-1:** MDNLCCFNSKFSKLAGGRLRGKQSRAYQVWLQPG*

**RG-pic3-2:** MDNLCCFNSKFSKLAGGRSSSLAKEEAIKGLPSMASAWLRGKLITPWKITMFLNLSNCMDMN*

**B**

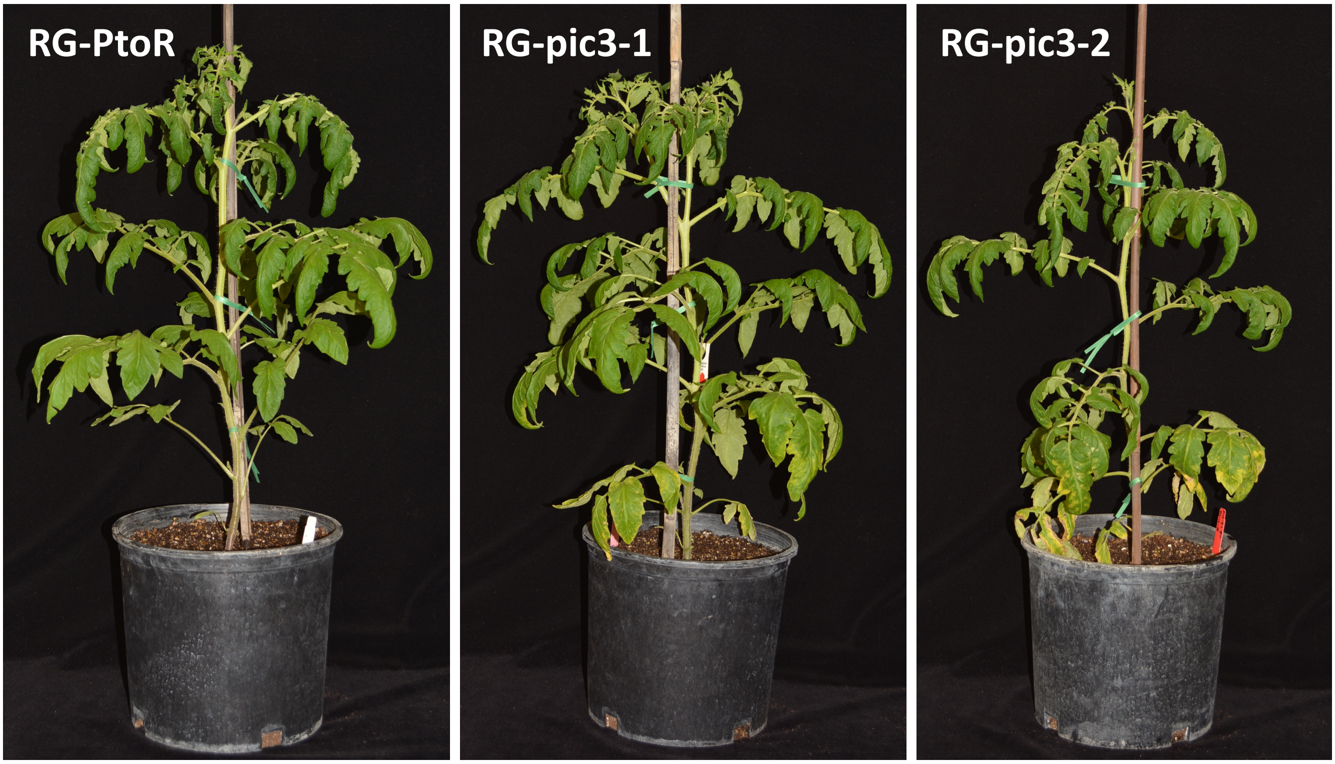


**Supplemental Figure S2. Possible truncated proteins produced in the RG-pic3 mutants and morphology of 8-week-old RG-pic3 mutant plants. A)** Pic3 protein sequence in RG-PtoR (wild type) and predicted truncated proteins that might be produced as the result of a premature stop codon in RG-pic3-1 and RG-pic3-2. The deletions in the RG-pic3-1 and RG-pic3-2 plants each cause a frame shift and introduce multiple amino acid substitutions around the cut site and eventually a premature stop codon at the 35$^{th}$ and 63$^{rd}$ amino acid of the Pic3 protein, respectively. Sequence highlighted in red in RG-PtoR indicates the PP2C domain predicted by NCBI Conserved Domains website. Asterisk "*" indicates the stop codon. **B)** Morphology of 8-week-old RG-PtoR, RG-pic3-1 and RG-pic3-2 plants.

Supplemental Figure S3

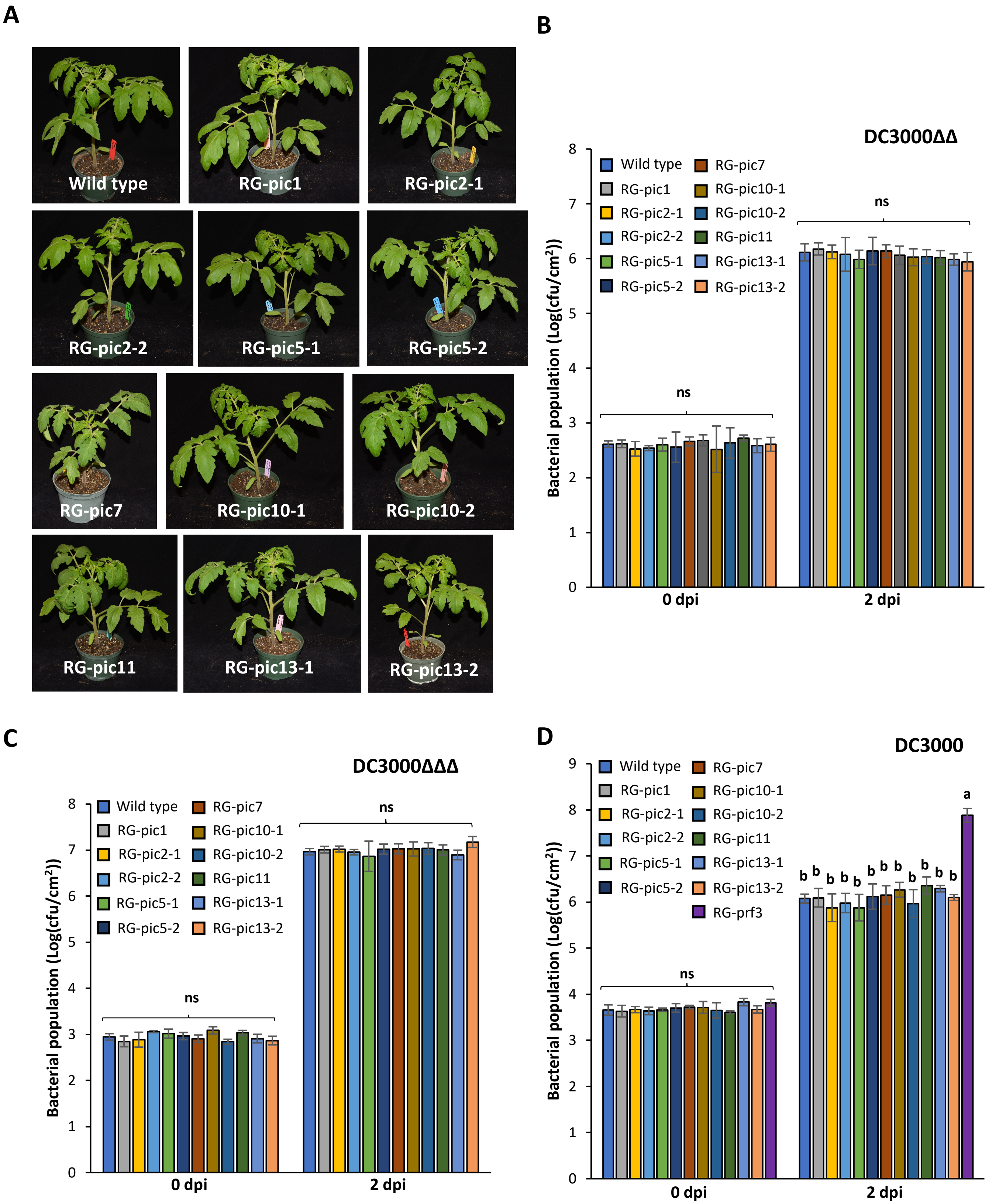

**Supplemental Figure S3. Loss-of-function mutations in seven tomato *Pic* genes do not affect resistance to *Pseudomonas syringae* pv. *tomato*. A)** Images of 4-week-old RG-PtoR (wild type) and multiple RG-pic mutant plants. **B)** RG-PtoR and RG-pic mutant plants were vacuum-infiltrated with $5\times10^4$ cfu/mL DC3000Δ*avrPto*Δ*avrPtoB* (DC3000ΔΔ). Bacterial populations were measured at 0 and 2 days post-inoculation (dpi). **C)** RG-PtoR and RG-pic mutant plants were vacuum-infiltrated with $5\times10^4$ cfu/mL DC3000Δ*avrPto*Δ*avrPtoB*Δ*fliC* (DC3000ΔΔΔ). Bacterial populations were measured at 0 and 2 dpi. **D)** RG-PtoR, RG-pic mutant and RG-prf3 plants were vacuum-infiltrated with $2\times10^5$ cfu/mL DC3000. Bacterial populations were measured at 0 and 2 dpi. Bars show means ± standard deviation, n = 3 or 4. cfu, colony-forming units. Different letters indicate significant differences based on a one-way ANOVA followed by Tukey's Honest Significant Difference post hoc test ($P < 0.05$). ANOVA was performed separately for each time point. ns, no significant difference. Graphs shown in (B, C and D) were made by combining results from separate small scale disease assays (i.e., from a few mutant plants). Bacterial populations of RG-PtoR from each assay were used for normalizing experimental data set. All inoculation experiments were repeated twice with similar results.

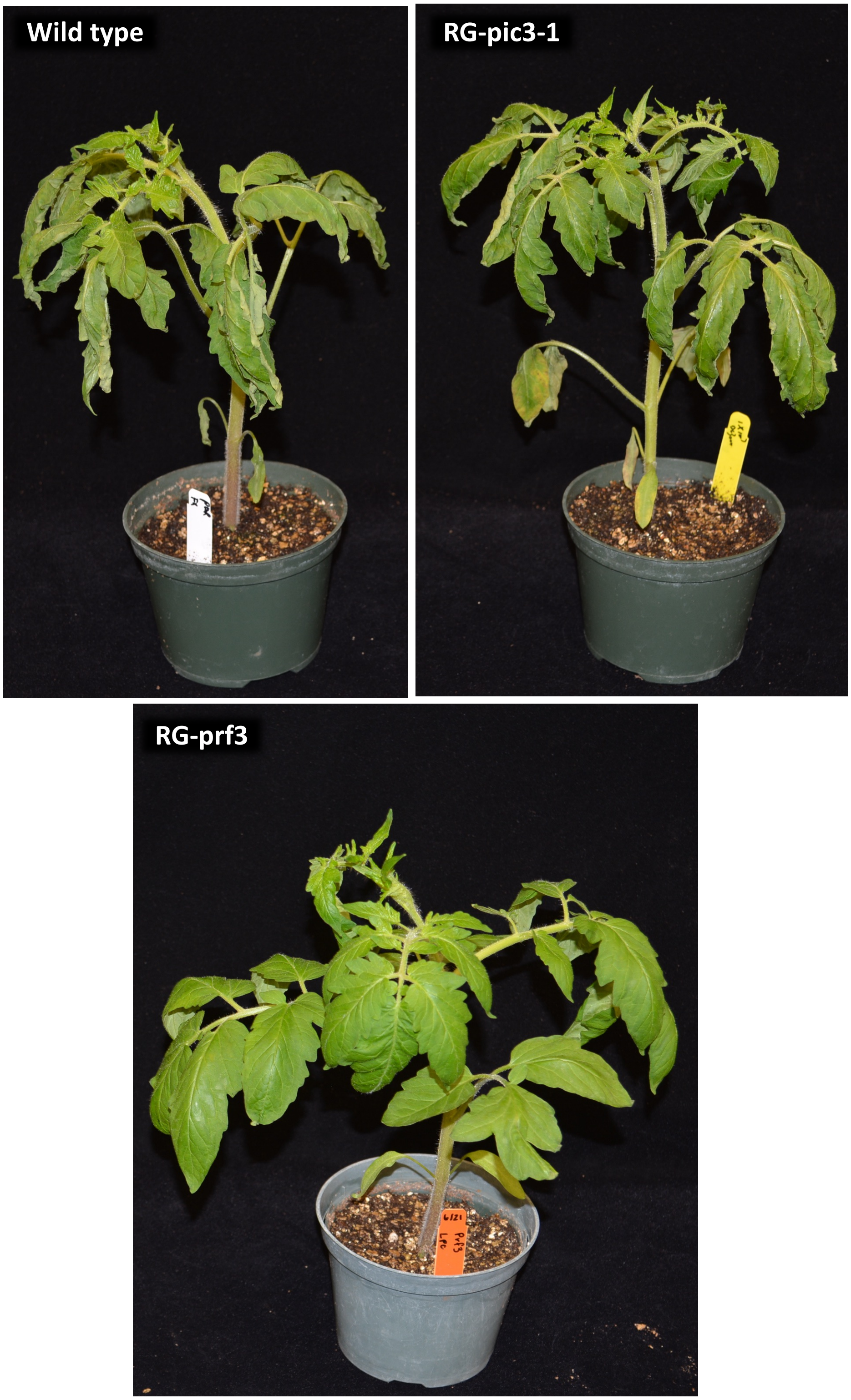


**Supplemental Figure S4. Loss-of-function mutation in *Pic3* does not affect the Pto/Prf3-mediated hypersensitive response (HR).** Wild-type (RG-PtoR), RG-pic3-1 and RG-prf3 plants (5-week-old) were vacuum-infiltrated with $1\times10^{7}$ cfu/mL DC3000 and photographed at 9 hours post-inoculation. The HR is seen as rapid leaf wilting in plants expressing Pto/Prf. RG-prf3 lacks Prf and does not develop an HR. Experiments were repeated twice with similar results.

### Supplemental Figure S5

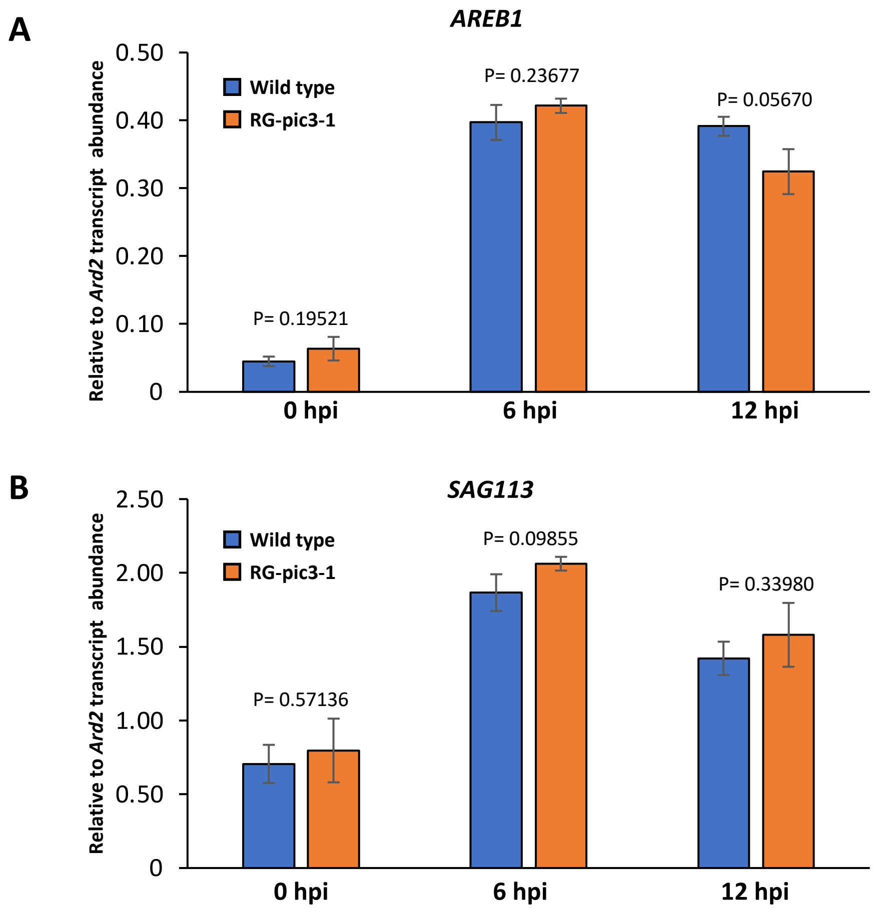


**Supplemental Figure S5. Transcript levels of ABA marker genes after ABA treatment.** Expression levels of *AREB1* (**A,** Solyc04g078840) and *SAG113* (**B**, Solyc05g052980) in older leaves of RG-PtoR (wild type) or RG-pic3-1 plants at 0, 6 and 12 h after 100 µM abscisic acid (ABA) syringe infiltration. Bars show means ± standard deviation, n = 3 plants. P values between wild type and RG-pic3-1 were calculated using Student's t-test. Experiments were repeated twice with similar results.

```
Pic2          DEISLKNSLDGIDLGNTETVVE--EISLENFLDGIDLGNTETVIDEISLENSLDGIDLGN   608
Pic1          EN-----------------FCSRSDTIFCGVFDGHGPY--GHMVARKVRDTLP-------   128
Pic13         PN-------------------LDSSTSFFGVYDGHGGD--E--VSKFCAKFLH-------   73
Pic10         AD--LAK----N---FGLNILGEDAISFYGVFDGHGGK--G--ASQFVRDYLP-------   146
Pic7          PL---------F---CKENSENSSNLHFFGVYDGHGCS--H--VAMKCKDRMH-------   165
Pic3          -S---------K---FV--QLHGHELGLFAIYDGHLGD--S--VPAYLQKHLF-------   87
Pic12         -A---------K---FM--QLRKGELGLFAIYDGHSGD--N--VAAYLQKHLF-------   85
Pic11         ------------------VISEEHGWVFVGIYDGFNGP--D--ATDFLLSNLY-------   303
AtPP2C38      QANNLLEDHSKLESGPVSMFDSGPQATFVGVYDGHGGP--E--AARFVNKHLF-------   104
Pic5          QANSLLEDQGQVF--------TTPSATYVGVYDGHGGP--Q--ASRFINNNLF-------   78
                                              *

Pic2          ----SSPMLHEFNLPIQIEKGDDPYQLL--EEYKIE-LDEGDIIVTATDALFDNLYDQEI   977
Pic1          F-----------------------GLISVPDVYYHRITDRNEFVVLATDGVWDVLSNKEA   345
Pic13         NKSLPA------------EK---QIVTANPDICTVELCNDDDFLVLACDGIWDCMSSQEV   283
Pic10         LKEVEK-------------G---GPLSAEPELKLLTLTKEDEFLIIGSDGIWDVFRSQNA   309
Pic7          ------------------------YVISEPEVTITDRTNEDECLILASDGLWDVVSNETA   341
Pic3          ------------------------HLSSDPDVTNADVDAETDLLILASDGLWKVMSNQEA   244
Pic12         ------------------------HLRSDPDVTTIDVDGDTDLLILASDGLWKVMSNQEA   242
Pic11         PKWNHA-LLEMFRIDYIGNS---PYINCLPSLHHHTLGSRDKFLILSSDGLYQYFTNEEA   553
AtPP2C38      AEFNREPLLAKFRVPEVFHK---PILRAEPAITVHKIHPEDQFLIFASDGLWEHLSNQEA   300
Pic5          PEFNRDPMFIQYGYPIPLKR---AVMSAEPSILIRKIRPEDLFLIFASDGLWDQLTDDEA   279
                                                              *
```

**Supplemental Figure S6. Sequence alignment of the tomato Pic and *Arabidopsis* PP2C38 proteins.** Full length sequence of Pic and *Arabidopsis* PP2C38 (At3g12620) proteins were used to conduct alignment, only the regions near the two conserved aspartic acid residues (highlighted in red) are shown. Pic7-05g (Solyc05g052980) and Pic10-07g (Solyc07g007220) were the source sequences of Pic7 and Pic10 used in this alignment analysis.

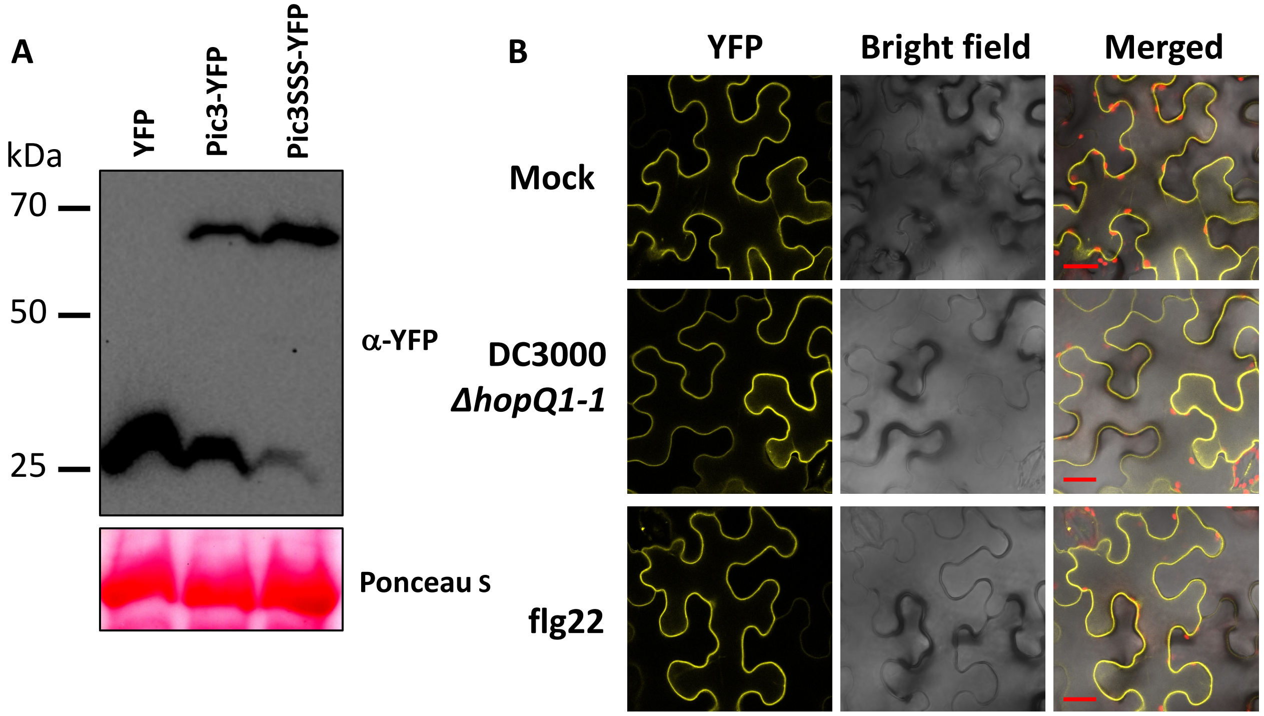


**Supplemental Figure S7. Pic3 protein subcellular localization is not altered by *Pseudomonas syringae* pv. *tomato* or flg22 treatment. A)** Western blotting analysis with an anti-YFP antibody confirmed that proteins tested in Figure 2E were expressed. *N. benthamiana* protein samples were collected at 40 h after initial Agrobacterium infiltration. **B)** Wild-type Pic3 proteins fused with YFP (pGWB541 vector) were transiently expressed in *N. benthamiana* leaves using Agrobacterium strain GV3101 for 40 h, the same Agrobacterium infiltrated spots were then syringe infiltrated with water, $1\times10^7$ cfu/mL DC3000*ΔhopQ1-1* or 1 µM flg22 peptide. After 1 h, the treated *N. benthamiana* leaves were examined under a confocal microscope. YFP fluorescence was excited by 485 nm laser. Scale bar, 25 µm. Experiments were repeated twice with similar results.

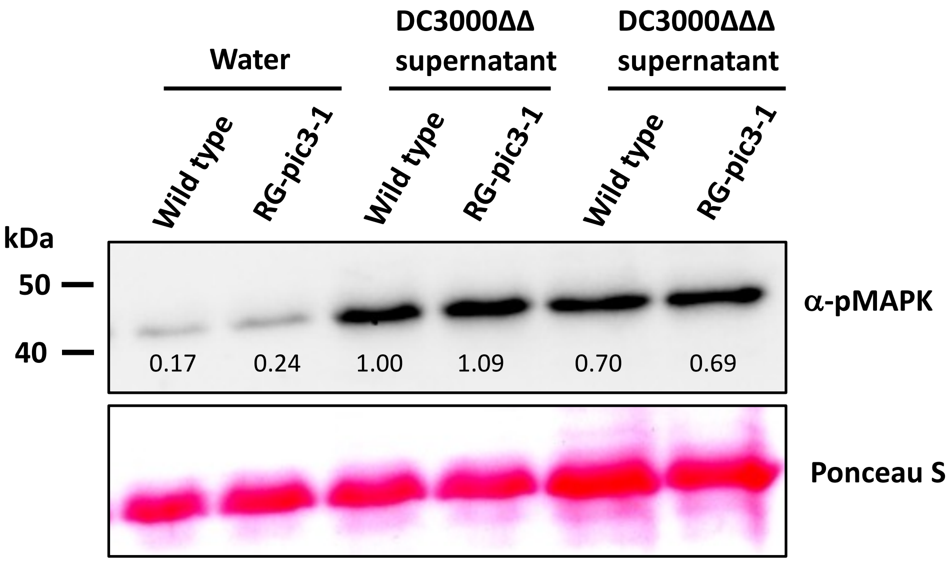


**Supplemental Figure S8. Tomato immunity-associated MAPK cascade is activated by a non-flagellin PAMP.** Immunoblot analysis with an anti-pMAPK antibody shows MAPK activation in RG-PtoR (wild type) and RG-pic3-1 plants by supernatants from boiled DC3000*ΔavrPtoΔavrPtoB* (DC3000ΔΔ) or DC3000*ΔavrPtoΔavrPtoBΔfliC* (DC3000ΔΔΔ). *P. s.* pv. *tomato* bacteria (1 mL of DC3000ΔΔ *or* DC3000ΔΔΔ, $OD_{600}$ = 1.0) were boiled at 95 °C for 20 min, followed by centrifugation at 10,000 g for 5 min to collect supernatant. Six leaf discs of 4-week-old plants (wild type (RG-PtoR) and RG-pic3-1) were floated in water overnight to let the wound response subside. The leaf discs were then incubated in 100× diluted supernatant or water (negative control) for 10 min and phosphorylated MAPK were detected with an anti-pMAPK antibody. Numbers indicated the phosphorylated MAPK protein levels quantified using ImageJ software.

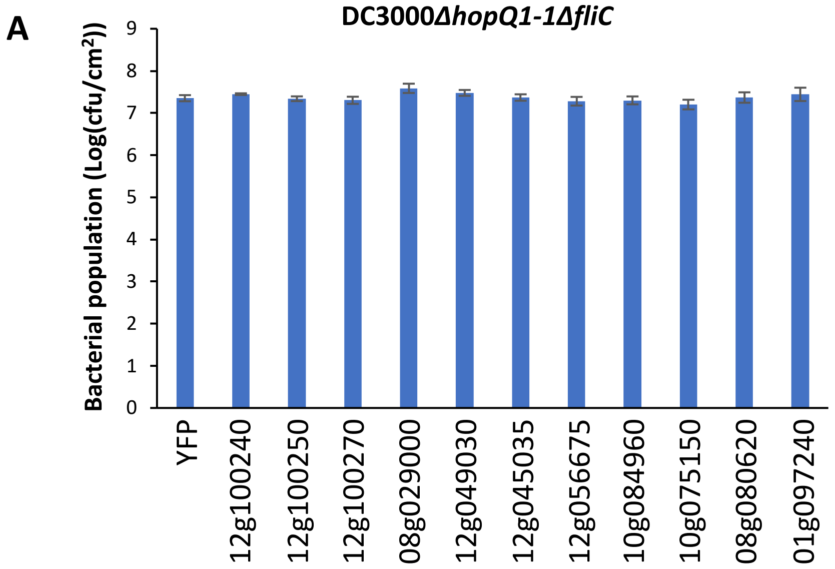


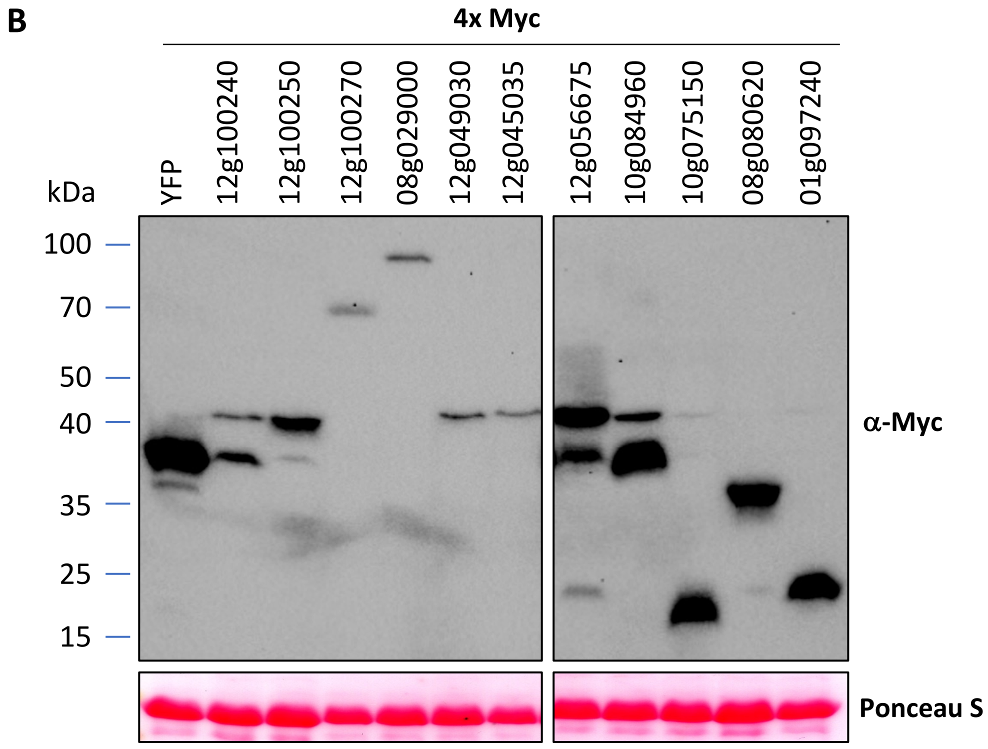


**Supplemental Figure S9. Expression of individual candidate genes did not alter *N. benthamiana* disease resistance. A)** Bacterial populations in *N. benthamiana* plants transiently expressing selective RNA-seq identified tomato or YFP (control) proteins 2 days after DC3000Δ*hopQ1-1*Δ*fliC* infiltration. Tomato (Solyc number provided) candidate or YFP protein was fused with Myc (4×) tag using pGWB417 vector. Bars show means ± standard deviation, n=4. No significant difference was detected between YFP and any of the selective RNA-seq identified proteins based on Student's t-test ($P < 0.05$). **B)** Immunoblot analysis confirmed that proteins were expressed in (A).

**Supplemental Figure S10**

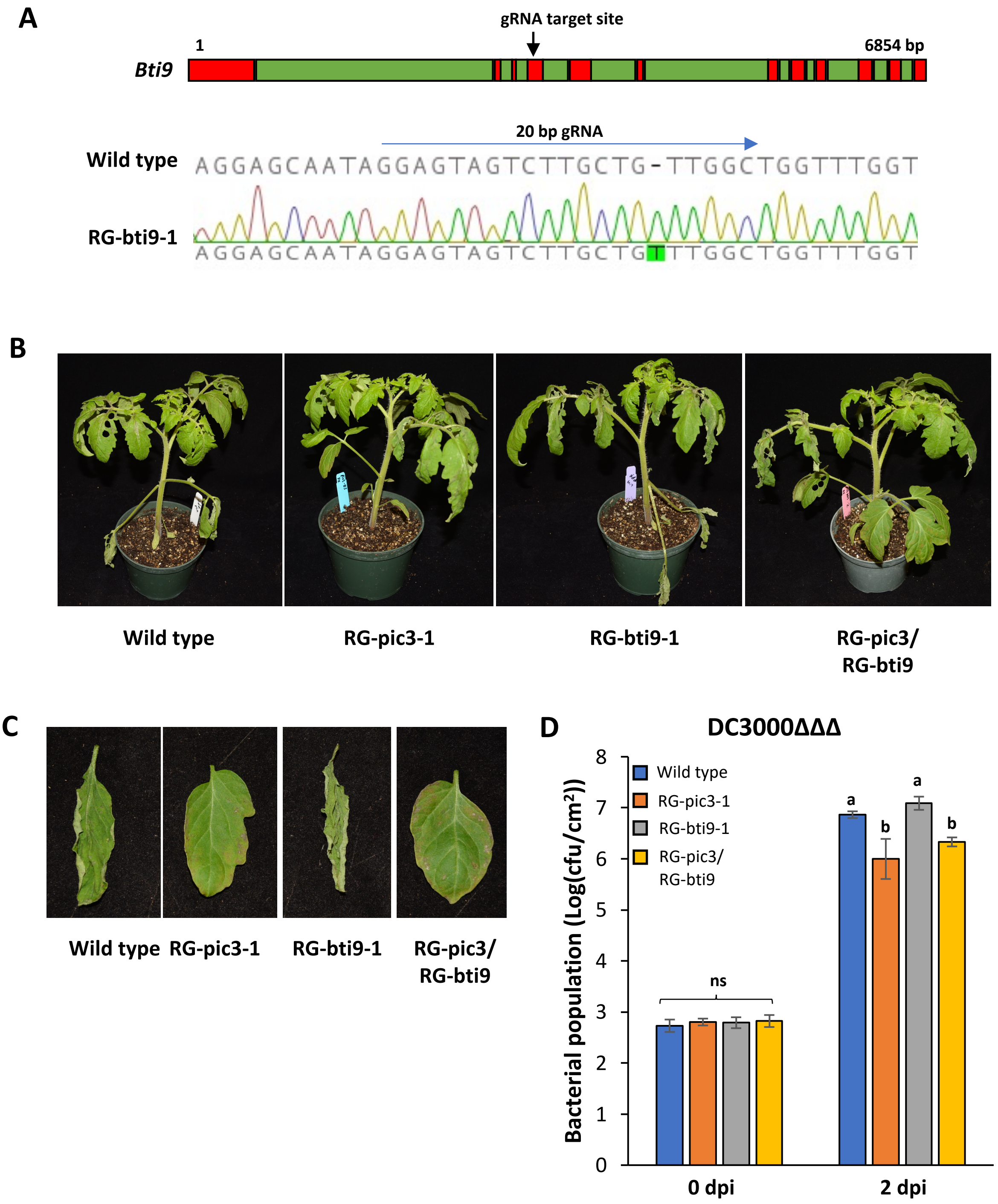


**Supplemental Figure S10. Loss-of-function mutation in *Bti9* does not abolish enhanced resistance in RG-pic3 mutant. A)** Top panel, schematic showing gene structure of *Bti9* (Solyc07g049180) including exons (red) and introns (green), arrow (black) points to CRISPR/Cas9 gRNA target site; Bottom panel, sequencing chromatogram of the RG-bti9-1 mutant allele (1 bp insertion in the fourth exon) generated by CRISPR/Cas9. **B)** Wild-type, RG-pic3-1, RG-bti9-1 and RG-pic3/RG-bti9 plants were vacuum-infiltrated with $5 \times 10^4$ cfu/mL DC3000Δ*avrPto*Δ*avrPtoB*Δ*fliC* (DC3000ΔΔΔ) and photographed at 3 days post-inoculation (dpi). **C)** Close-up images of the older leaves from (B). **D)** Bacterial populations in older leaves (1st or 2nd true leaf) were measured at 0 and 2 dpi. Bars show means ± standard deviation, n = 3 or 4. cfu, colony-forming units. Different letters indicate significant differences based on a one-way ANOVA followed by Tukey's Honest Significant Difference post hoc test ($P < 0.05$). ANOVA was performed separately for each time point. ns, no significant difference. Experiments were repeated three times with similar results.

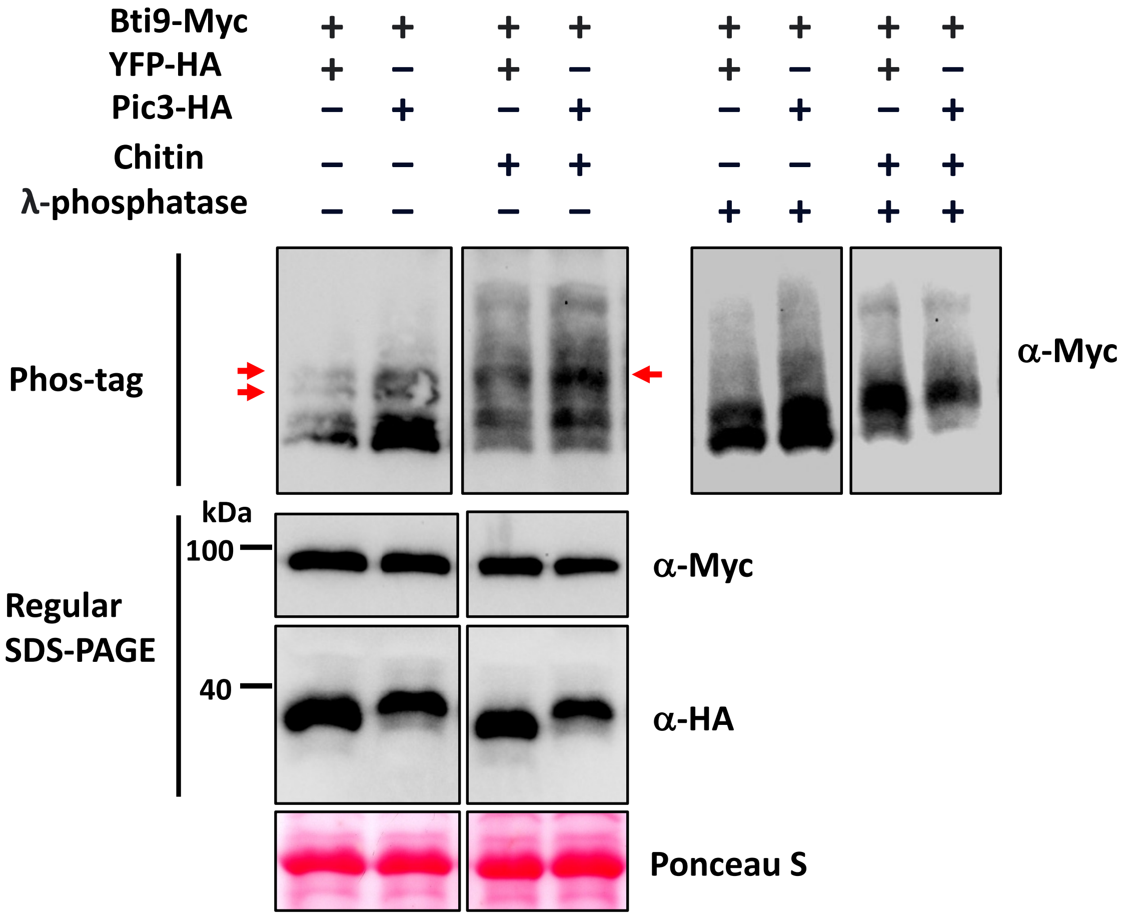


**Supplemental Figure S11. Pic3 does not appear to affect Bti9 phosphorylation status.** Bti9, YFP or Pic3 proteins (fused with epitope tags, as shown) were transiently expressed using agroinfiltration of leaves of *N. benthamiana* with Agrobacterium strain GV2260. After 40 h, the same agroinfiltrated spots were syringe-infiltrated with either water or 200 µg/mL chitin. After 30 min, leaf samples were collected for protein extraction. Protein extracts were either separated on 12% SDS-PAGE gel and detected by an anti-Myc or anti-HA antibody or separated on 8% SDS-PAGE gel containing 40 µM Phos-tag Acrylamide AAL107 (FUJIFILM Wako Chemicals) and detected by an anti-Myc antibody. Protein extracts were treated with λ-phosphatase (NEB Lambda protein phosphatase, P0753S) following manufacturer's instructions. Arrows show the main phosphorylated proteins that were affected by λ-phosphatase. Experiment was repeated twice with similar results.

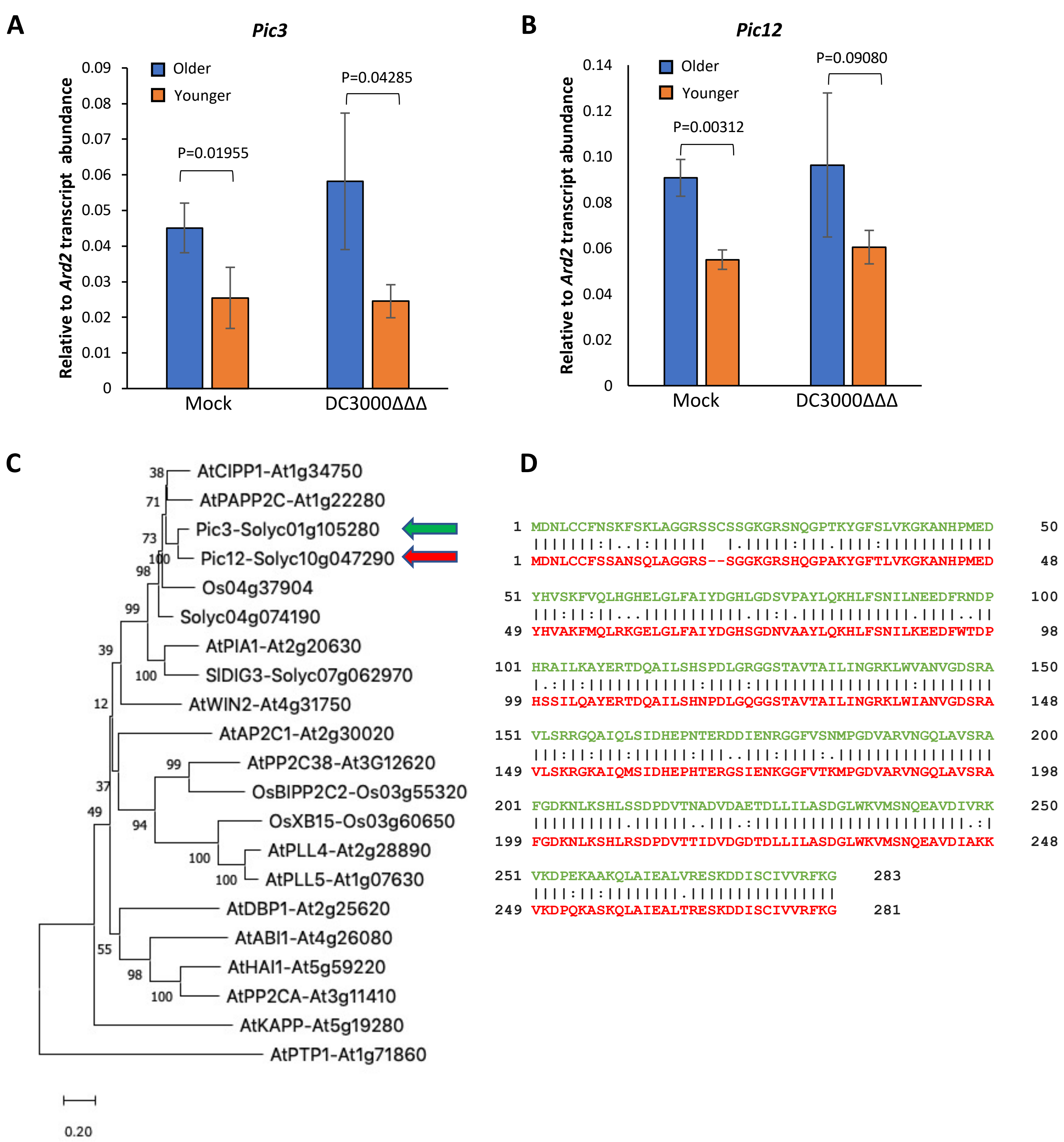


**Supplemental Figure S12. Pic12 and Pic3 are closely related PP2C phosphatases. A)** *Pic3* transcript abundance in leaves of RG-PtoR (wild type) plants at 6 h after mock (10 mM $MgCl_2$) or 5 $\times 10^6$ cfu/mL DC3000*ΔavrPtoΔavrPtoBΔfliC* (DC3000ΔΔΔ) vacuum infiltration. **B)** *Pic12* transcript abundance in leaves of RG-PtoR (wild type) plants at 6 h after mock (10 mM $MgCl_2$) or 5 $\times 10^6$ cfu/mL DC3000ΔΔΔ vacuum infiltration. 'Older' refers to $1^{st}$ or $2^{nd}$ true leaf and 'younger' refers to $4^{th}$ or $5^{th}$ true leaf. Bars show means ± standard deviation, n = 3 plants. P values between older and younger were calculated via Student's t-test. No significant difference between mock and DC3000ΔΔΔ was detected for either older or younger leaves based on Student's t-test ($P < 0.05$). **C)** Phylogenetic tree constructed using amino acid sequences of Pic3, Pic12 and related PP2C phosphatases in *Arabidopsis*, tomato and rice. Clustalx2.1 was used to conduct sequence alignment and MEGA11 was used to construct the neighbor joining phylogenetic tree, bootstrap was set as 1000. **D)** Amino acid sequence alignment between Pic3 (green) and Pic12 (red).

**A** **Supplemental Figure S13**

**RG-PtoR:** MDNLCCFSSANSQLAGGRSSGGKGRSHQGPAKYGFTLVKGKANHPMEDYHVAKFMQLRK
GELGLFAIYDGHSGDNVAAYLQKHLFSNILKEEDFWTDPHSSILQAYERTDQAILSHNPDLGQ
GGSTAVTAILINGRKLWIANVGDSRAVLSKRGKAIQMSIDHEPHTERGSIENKGGFVTKMPG
DVARVNGQLAVSRAFGDKNLKSHLRSDPDVTTIDVDGDTDLLILASDGLWKVMSNQEAVDI
AKKVKDPQKASKQLAIEALTRESKDDISCIVVRFKG*

**RG-pic12-1:** MDNLCCFSSANSQLAGGRSSGGKGRSHQGPAKYGFTLVKGKGGLSCC*

**RG-pic12-2:** MDNLCCFSSANSQLAGGRSSGGKGRSHQGPAKYGFTLVKGKANHPTGGLSCC*

**B**

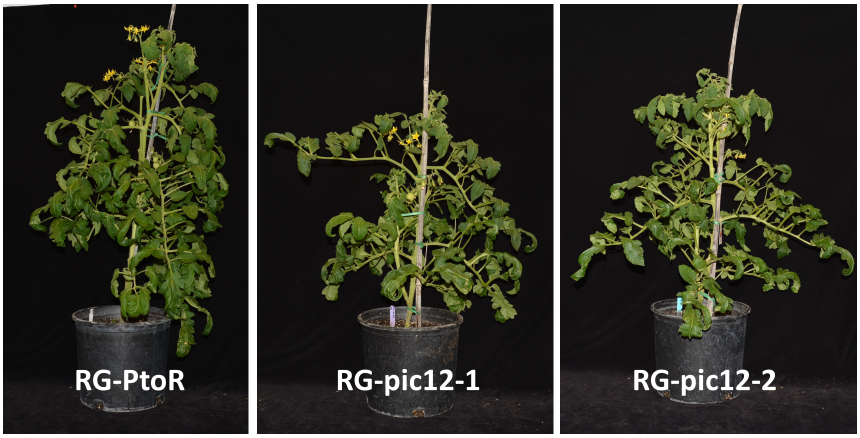


**C**

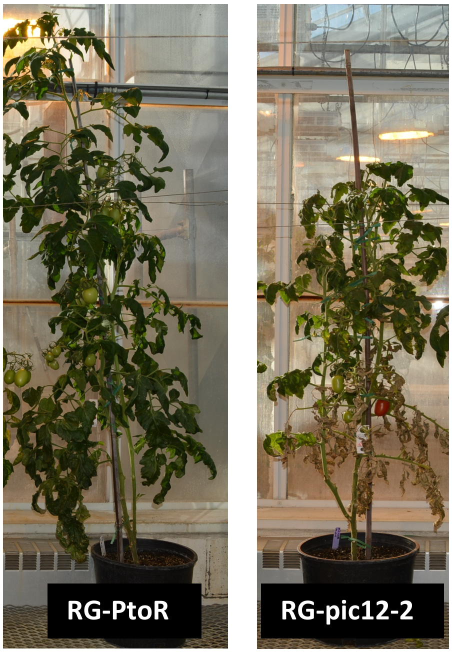


**Supplemental Figure S13. Predicted truncated proteins produced in the RG-pic12 mutants and morphology of 8-week-old wild type and RG-pic12 plants. A)** Pic12 amino acid sequences in RG-PtoR (wild type) and possible truncated proteins that could be produced as the result of a premature stop codon in RG-pic12-1 and RG-pic12-2. The deletions and insertions in the RG-pic12-1 and RG-pic12-2 lines cause a frameshift and introduce multiple amino acid substitutions around the cut site and eventually a premature stop codon at the 48$^{th}$ and 53$^{rd}$ amino acid of the Pic12 protein, respectively. Sequence highlighted in red in RG-PtoR indicates the PP2C domain predicted by NCBI Conserved Domains website. Asterisk "*" indicates the stop codon. **B)** Morphology of 8-week-old RG-PtoR, RG-pic12-1 and RG-pic12-2 plants. **C)** Morphology of 12-week-old RG-PtoR and RG-pic12-2 plants.

Supplemental Figure S14

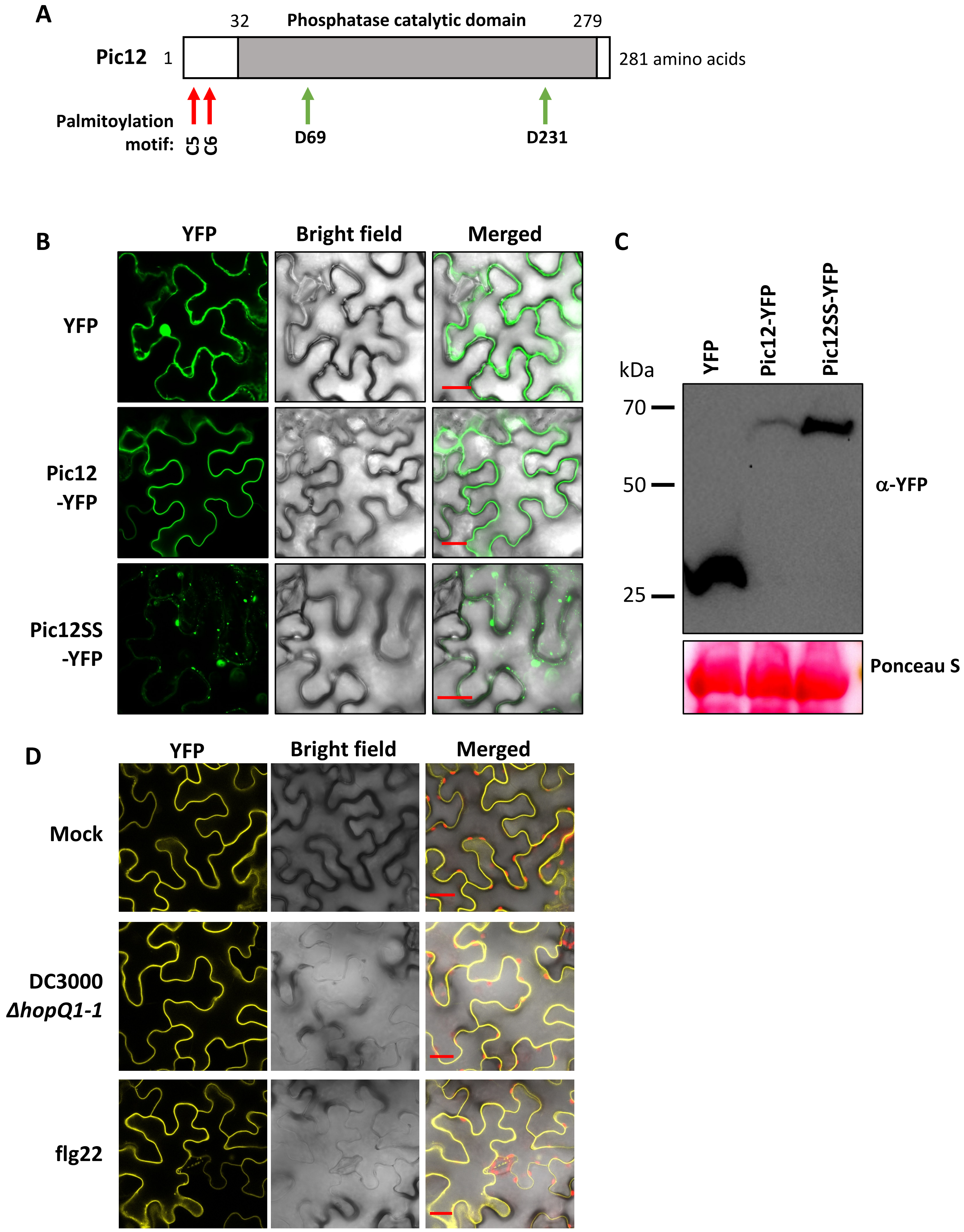

**Supplemental Figure S14. Palmitoylation motif affects Pic12 protein subcellular localization. A)** Schematic of Pic12 protein showing the two predicted palmitoylation sites (red arrows) at the N terminus and two conserved aspartic acid residues (green arrows) involved in metal binding in the PP2C catalytic domain. **B)** Wild-type Pic12 or the Pic12SS variant protein fused with YFP (pGWB541 vector) were transiently expressed in *N. benthamiana* leaves using Agrobacterium strain GV3101 for 40 h and examined under a confocal microscope. YFP fluorescence was excited by 485 nm laser. The pGWB541 vector (CmR, ccdB removed) expressing YFP was used as a control. Scale bar, 25 µm. Experiments were repeated twice with similar results. **C)** Western blotting analysis with an anti-YFP antibody confirmed that proteins tested in (B) were expressed. *N. benthamiana* protein samples were collected at 40 h after initial agrobacterium infiltration. **D)** Wild-type Pic12 protein fused with YFP (pGWB541 vector) was transiently expressed in *N. benthamiana* leaves using Agrobacterium strain GV3101 for 40 h. The same agrobacterium infiltrated spots were then syringe infiltrated with water, $1\times10^7$ cfu/mL DC3000*ΔhopQ1-1* or 1 µM flg22 peptide. After 1 h, the treated *N. benthamiana* leaves were examined under a confocal microscope. YFP fluorescence was excited by 485 nm laser. Scale bar, 25 µm. Experiments were repeated twice with similar results.

**Supplemental Figure S15**

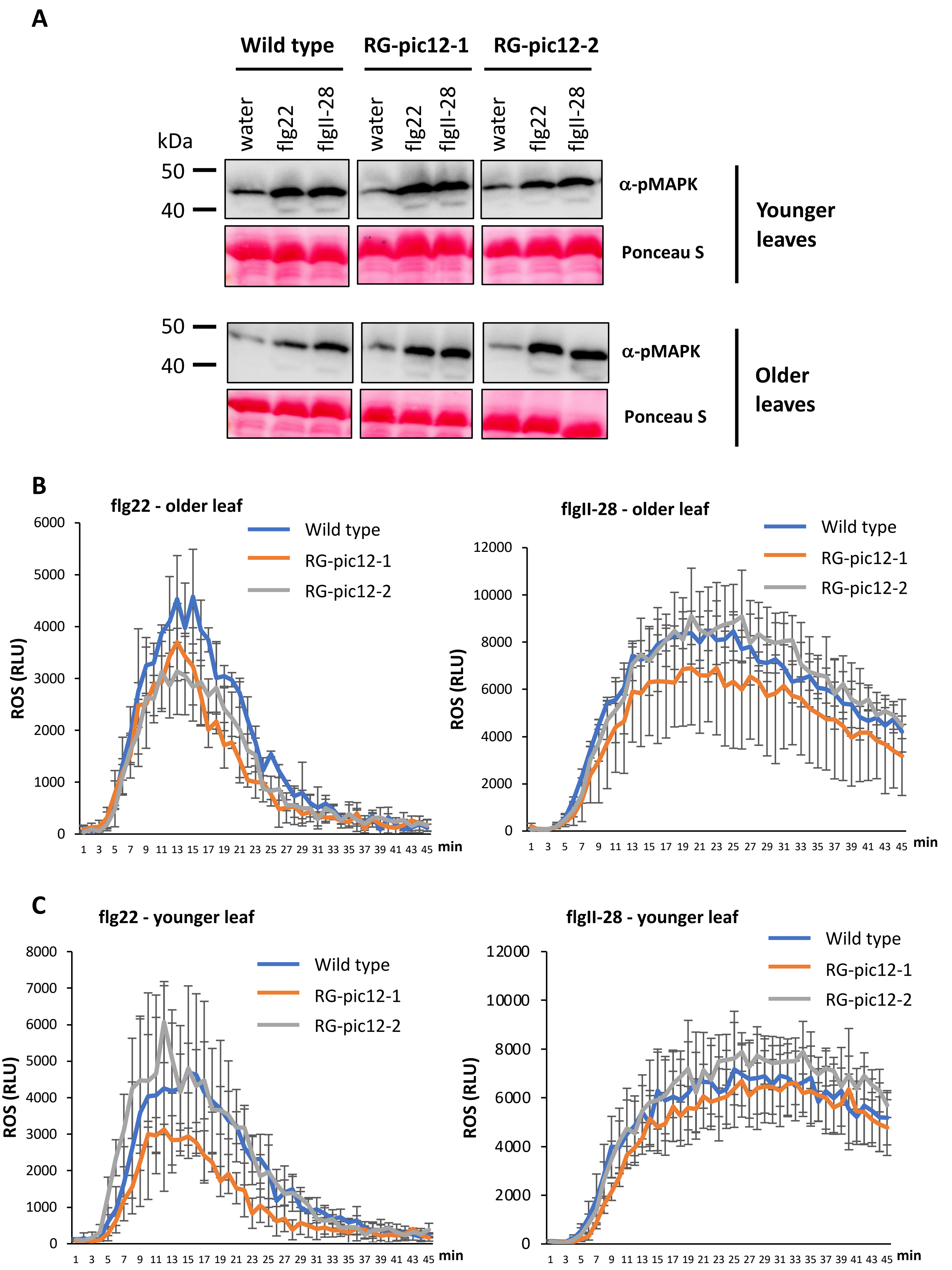

**Supplemental Figure S15. Loss-of-function mutations in *Pic12* do not affect MAPK activation or ROS production in response to flg22 and flgII-28. A)** MAPK activation in wild-type (RG-PtoR) and RG-pic12 mutant plants. Water, 10 nM flg22 or 25 nM flgII-28 were applied to detached leaf discs for 10 min and proteins were separated on 12% SDS-PAGE gel and detected by an anti-pMAPK antibody. **B** and **C)** Time course ROS production assay in wild-type and RG-pic12 mutant plants. 100 nM flg22 or 100 nM flgII-28 were used to activate ROS production in detached leaf discs and relative light units (RLUs) were measured for 45 min. Bars show means ± standard deviation, n = 3 plants. No significant differences were detected between wild-type and RG-pic12 plants at peak readout based on a one-way ANOVA followed by Tukey's Honest Significant Difference post hoc test ($P < 0.05$). Experiments were repeated three times with similar results.

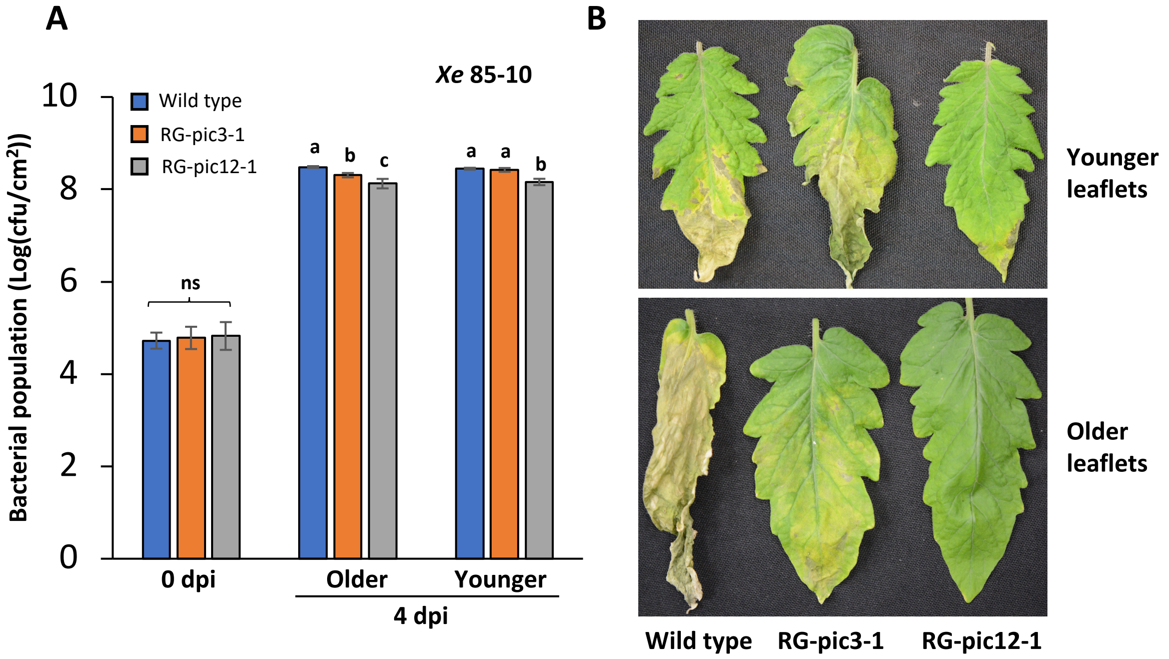


**Supplemental Figure S16. Loss-of-function RG-pic3 and RG-pic12 mutants show enhanced disease resistance to *Xanthomonas euvesicatoria*. A)** Wild-type (RG-PtoR), RG-pic3-1 and RG-pic12-1 plants were vacuum-infiltrated with $2 \times 10^5$ cfu/mL *Xanthomonas euvesicatoria* strain 85-10. Bacterial populations were measured at 0 and 4 days post-inoculation (dpi). 'Older' refers to $2^{nd}$ true leaf and 'younger' refers to $5^{th}$ true leaf. Bars show means ± standard deviation, n = 3. cfu, colony-forming units. Different letters indicate significant differences based on a one-way ANOVA followed by Tukey's Honest Significant Difference post hoc test ($P < 0.05$). ANOVA was performed separately for each time point and leaf type. ns, no significant difference. Experiments were repeated three times with similar results. **B)** Disease symptoms on younger and older leaves of indicated genotypes related to experiment in (A), photographed at 8 dpi.

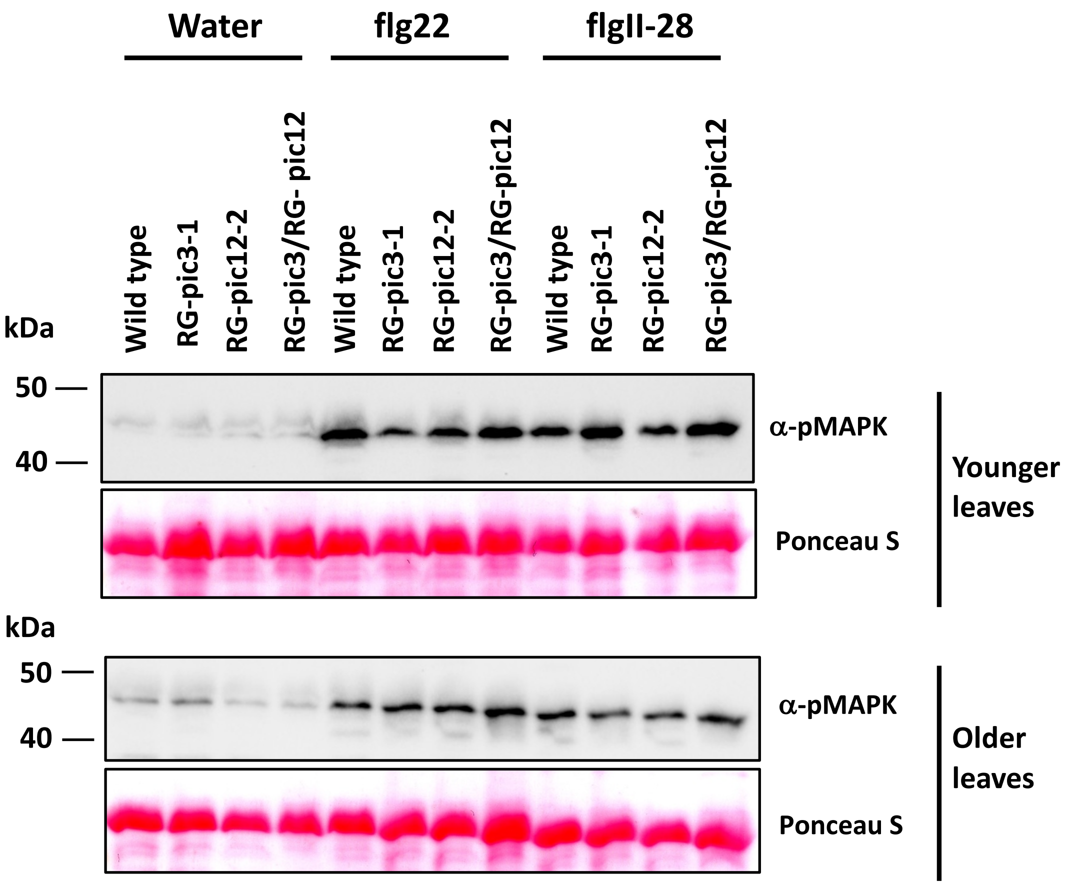


**Supplemental Figure S17. Loss-of-function mutations in both *Pic3* and *Pic12* do not affect MAPK activation in response to flg22 and flgII-28.** MAPK activation in wild-type, RG-pic3-1, RG-pic12-2 and RG-pic3/RG-pic12 mutant plants. Water, 10 nM flg22 or 25 nM flgII-28 were applied to detached leaf discs for 10 min and proteins were separated on 12% SDS-PAGE gel and detected by an anti-pMAPK antibody. 'Older' refers to 1$^{st}$ or 2$^{nd}$ true leaf and 'younger' refers to 4$^{th}$ or 5$^{th}$ true leaf. Experiments were repeated three times with similar results.
